# Soil Chemistry and Microbiome Determine N_2_O Emission Potential in Soils

**DOI:** 10.1101/2020.12.16.422796

**Authors:** Matthew P. Highton, Lars R. Bakken, Peter Dörsch, Sven Tobias-Hunefeldt, Lars Molstad, Sergio E. Morales

## Abstract

Microbial nitrogen (N) transformations in soil, notably denitrification, result in the production of the potent greenhouse and ozone depleting gas nitrous oxide (N_2_O). Soil chemistry and microbiome composition impact N_2_O emission potential but the relative importance of these factors as determinants of N_2_O emission in denitrifying systems is rarely tested. In addition, previous linkages between microbiome composition and N_2_O emission potential rarely demonstrate causality. Here, we determined the relative impact of microbiome composition (i.e. soil extracted cells) and chemistry (i.e. water extractable chemicals) on N_2_O emission potential utilizing an anoxic cell based assay system. Cells and chemistry for assays were sourced from soils with contrasting N_2_O/N_2_O+N_2_ ratios, combined in various combinations and denitrification gas production was measured in response to nitrate addition. Average directionless effects of cell and chemical extract on N_2_O/N_2_O+N_2_ (Cell: Δ0.16, Chemical extract: Δ0.22) and total N_2_O hypothetically emitted (Cell: Δ2.62 μmol-N, Chemical extract: Δ4.14 μmol-N) indicated chemistry is the most important determinant of N_2_O emissions. Independent pH differences of just 0.6 points impacted N_2_O/N_2_O+N_2_ on par with independent chemical extract differences, supporting the dominance of this variable in previous studies. However, impacts on overall N_2_O hypothetically emitted were smaller suggesting that soil pH manipulation may not necessarily be a successful approach to mitigate emissions over a fixed time period. In addition, we observed increased N_2_O accumulation and emission potential at the end of incubations concomitant with predicted decreases in carbon availability suggesting that carbon limitation increases N_2_O emission transiently with the magnitude of emission dependent on the both chemical and microbiome controls.

## Introduction

Nitrous oxide (N_2_O) is a potent greenhouse gas and ozone depleter accounting for around 6.2 percent of worldwide greenhouse gas emissions on a CO_2_ mass equivalence basis (Intergovernmental Panel on Climate Change, 2013). Around 45% percent of this is anthropogenically produced, mostly (60%) in agricultural settings via soil based N transformations (Syakila and Kroeze, 2011). Denitrification, the anaerobic microbial reduction of N compounds (NO_3_^−^➔NO_2_^−^➔NO➔N_2_O➔N_2_), is considered a major pathway of anthropogenic N_2_O production (Bouwman *et al.*, 2013). Soil conditions (e.g. O_2_ concentration (Zumft, 1997; Smith and Tiedje, 1979; Firestone *et al.*, 1979) and pH (Simek and Cooper, 2002; Liu *et al.*, 2014; Čuhel and Šimek, 2011)), can affect the ratio of the major gaseous end products of this process (N_2_O & N_2_) and overall process rates resulting in higher or lower N_2_O emissions to the atmosphere. Therefore, understanding the soil factors that favour low N_2_O emission in the presence of available soil N is of great importance to manipulating agricultural systems towards reduced N_2_O production in the future.

Conceptually, factors affecting soil N_2_O emission potential can be separated into three areas: distal controls which act in the long term to determine denitrifier microbiome composition, the genetic and regulatory potential of the microbiome itself, and the immediate scale impact of proximal controls which may be transduced through the denitrifiers present (Wallenstein *et al.*, 2006). Proximal factors such as O_2_, pH and temperature are easily isolated as independent variables, making them ideal experimental targets. In contrast, microbiome impacts are difficult to isolate due to confounding by soil chemical and physical factors, likely distal controls. As such, they are more poorly understood. Studies are often suggestive (Graf *et al.*, 2014) or correlative (Samad *et al.*, 2016; Jones *et al.*, 2014; Philippot *et al.*, 2009; Čuhel *et al.*, 2010; Morales *et al.*, 2010) and it is often unclear whether microbiome features described are the true driver of an N_2_O emission outcome. The issue is exacerbated when co-variance is observed with variables such as pH, which are known to affect both N_2_O/N_2_ emission ratios and changes in microbiome composition (Samad *et al.*, 2016; Philippot *et al.*, 2009).

Attempts have been made to control “all” variables relevant to denitrification within soils to isolate microbiome based effects, however, this may not account for the effect of physical differences between the soils and certainly doesn’t for unknown and unaccounted variables impacting denitrification gas kinetics at the time of experimentation (Cavigelli and Robertson, 2000; Holtan-Hartwig *et al.*, 2000). A solution to such problems may be the extraction of whole microbiomes from soils. Though probably biased in the portion of soil communities extracted e.g. (Nadeem *et al.*, 2013; Holmsgaard *et al.*, 2011), this method has demonstrated that communities from different soils or the same soil under different long term pH treatments will show contrasting N_2_O emission responses to the same pH conditions (Dörsch *et al.*, 2012; Liu *et al.*, 2014).

Despite an increasing focus on microbiome impacts, the relative impact of proximal effects vs. microbiome composition on N_2_O emission from denitrification is still poorly understood. In practice, should management of soil to control N_2_O emissions be targeted towards proximal effects, or is the long term selection of certain denitrifier community biomes (distal control) more important?

Here, we incubated soil extracted cells in chemical extracts from pairs of soils with contrasting N_2_O/N_2_O+N_2_ emission ratios in all potential combinations with the aim of identifying whether microbiome composition (extracted cell origin) or proximal control (extracted chemical environment) in general was the most important determinant of the contrasting N_2_O/N_2_O+N_2_ ratios and total N_2_O emission in our model system and soils in general. We hypothesized chemical differences (especially pH) would be the dominant effector while microbiome composition effects would weaker but still detectable. Soil cell extraction allowed treatment of microbial communities as independent transferable units while extraction of soil chemistry ensured that whatever water-extractable components of the soil were present (e.g. dissolved carbon) reflected the parent soil in the produced incubation media. This is in contrast to traditional lab-based analyses which typically use a single simple carbon source.

## Methods

### 2.1 Soil sampling

Soils were re-sampled from New Zealand South Island pasture farms (Karangarua, Makarora, Tapawera, Fairlie-Geraldine, Woodend, Rae’s Junction) previously sampled in Highton et al. (2020). Sampling took place from 21^st^ to 23^rd^ of March, 2018. Soils were selected based on contrasting pH and N_2_O hypothetically emitted (%) identified in Highton et al. (2020). Multiple soil cores (10cm length, 2.5cm diameter) were sampled along a 7.5m transect evenly at distances of 0, 2.5, 5 and 7.5m using a foot-operated auger until ~3kg of soil was collected. Repeated cores at each distance were carried out in 4 perpendicular rows up to 6 cores across. Pooled site cores were stored field moist on ice in partially open ziplock bags during transport and at 4ºC in the lab. Grass, insects, worms and large roots were removed and cores were sieved at 2mm. Sieved soils were stirred rigorously with a metal spoon to homogenize. Soils underwent a 36hr period without temperature control during transport to the Norwegian University of Life Sciences (NMBU, Ås, Akershus, Norway).

### 2.2 Soil pH

Soil pH was measured using both CaCl_2_ (10mM) and ddH_2_O extractants as in Highton et al. (2020). Values were measured using an Orion 2 star pH meter (ThermoFisher Scientific, Waltham, Massachusetts, USA) with an Orion Ross Sure Flow Electrode (ThermoFisher Scientific), allowing up to 5 minutes for readings to stabilize.

### 2.3 Anoxic soil incubations

Anoxic soil incubations were carried out to determine soil denitrification gas kinetics and N_2_O emission potential. Incubations were prepared as in Highton et al. (2020) excluding overnight storage and oxic preincubation. Briefly, 3mM NH_4_NO_3_ was amended to soils by a flooding and draining procedure. Twenty grams dry weight equivalent of soil were weighed into triplicate 120ml serum vials per soil. Vials were crimp sealed with butyl rubber septa and made anoxic by repeated evacuation and helium flushing, Soil vials were incubated at 20ºC in a temperature controlled water bath. Headspace gases (1ml) were sampled every 4hrs via an automated robotic gas sampling system (Molstad *et al.*, 2007, 2016). Gases (O_2_, CO_2_, NO, N_2_O and N_2_) were quantified in real time using a coupled Agilent 7890A gas chromatograph (GC) equipped with an ECD, TCD, FID, and chemiluminescence NOx analyser (Model 200A, Advanced Pollution Instrumentation, San Diego, USA). An equal volume of helium is returned to the vials by back pumping ensuring consistent vial pressure. Dilution of headspace gases is accounted for later through back calculation. Gas concentrations were calibrated using premixed standard gases supplied by AGA industrial gases (Oslo, Akershus, Norway). The overall system and its improvements are described in detail in (Molstad *et al.*, 2007, 2016).

### 2.4 Cell based assay

A soil extracted cell based assay (CBA) was developed to determine the relative importance of microbiome composition and soil chemistry on N_2_O emission potential (see emission potential metrics 2.7). Extraction of soil components allowed them to be treated as independent experimental units. Soil chemistry and cells were extracted separately from soils with similar native pH and contrasting N_2_O emission potential: Karangarua, a low N_2_O emitting soil (N_2_O hypo emit ratio = 0.26, pH = 5.75) and Rae’s Junction, a high N_2_O emitting soil (N_2_O hypo emit ratio = 0.92, pH = 5.6). Extracted cells and chemistry were combined in 4 possible combinations to give the standard treatments: High emitting cells (HEC) + high emitting extract (HEE), high emitting cells (HEC) + low emitting extract (LEE), low emitting cells (LEC) + high emitting extract (HEE), low emitting cells (LEC) + low emitting extract (LEE). Standard treatments were carried out in triplicate vials. Minimum duplicate 3mM glutamate amended controls of each treatment were produced to understand the impact of carbon limitation. Duplicate chemical extract free control incubations containing just extracted cells and milliQ were prepared to test the baseline activity of extracted cells. Occasional replication in duplicate was necessitated by limited vial space in the automated incubator/gas sampler. Cell negative controls were prepared to confirm the sterility of chemical extracts and to quantify the elution of any N_2_ and O_2_ remaining in the extract media after He flushing. Full treatment contents and replication is detailed in Table S1. Hereafter this initial cell based assay is referred to as CBA-int to differentiate it from the CBA using alternate pH soils (section 2.5)

#### 2.4.1 Chemical extract media preparation

Water extractable organic carbon (WEOC) extraction was based on a previous protocol (Guigue *et al.*, 2014). Air-dried soil was combined with milliQ H_2_O at a 1:3 ratio (170g: 510ml) in 1L Schott bottles. Bottles were shaken lengthways on an orbital shaker at 120rpm for 1hr. Coarse particles were allowed to settle out for 5 minutes and supernatant was poured into 250ml polycarbonate Nalgene centrifuge tubes (ThermoFisher). Fine particles were removed by successive centrifugation (pelleting) and filtration steps: centrifugation at 4600G for 20minutes using JXN-26 high-speed centrifuge with JS-7.5 swing out rotor (Beckman Coulter, Brea, California, USA), filtration using 500ml Sterafil Filter Holders (Merck, Burlington, MA, USA) loaded with 1.2μm glass-fibre pre filters (Merck) and 0.45μM cellulose filters (Merck), syringe filtration using sterile 0.22μm mixed cellulose ester filters (Merck). Filter sterilized Na-glutamate solution was added to a portion of the chemical extract solution from each soil to give a final concentration of 3mM once diluted in final treatment vials. An equivalent volume of milliQ H_2_O was added to the rest of the extract to account for dilution. Standard extracts, glutamate amended extracts and milliQ for carbon free controls were buffered to pH 6 using 20mM Na-phosphate buffer, as this was the closest value to parent soil pH H_2_O (Rae’s Junction= 5.60, Karangarua = 5.75) within the bufferable range. Extracts and milliQ were re-filtered at 0.22μm to ensure sterility after pH and carbon manipulation. 22.5ml of solution was added to autoclaved 120ml glass serum vials containing magnetic stir bars. Vials were crimp sealed with butyl rubber septa + aluminium cap. Anoxia was induced through 8 repeated cycles of vacuum evacuation and helium filling with continuous magnetic stirring at 360rpm. Vials were stored at 8ºC until inoculation and incubation.

#### 2.4.2 Cell extraction by low speed centrifugation

The cell extraction procedure was modified from (Lindahl and Bakken, 1995) with cell separation on the basis of sedimentation rate using low speed centrifugation. Cell extractions were performed on the same day they would be used, using optimized conditions determined in an earlier test extraction yielding approximate cell extraction efficiencies for each soil. Twenty g of field moist soil was blended with 200ml of milliQ H_2_O in a two speed Waring blender (Waring, Stamford, Connecticut, USA) on high for 3×1min with 5min intermittent cooling on ice between each blending run. Coarse particles were allowed to settle for 5min before supernatant was poured off into sterile falcon tubes up to the 35ml mark (equivalent to 8cm centrifugation distance). Tubes were centrifuged at 1000G for 10minutes with 4ºC cooling on a benchtop Mega star 1.6R centrifuge with a TX-150 swing out rotor (VWR, Radnor, Pennsylvania, US) to sediment out non-cellular debris. Cell containing supernatant was recovered into additional falcon tubes and centrifuged at 10,000G for 20 minutes with 4ºC cooling to pellet cells using an Avanti JXN-30 highspeed centrifuge with JA 14.50 fixed angle rotor (Beckman Coulter). Supernatant was removed without disturbing the cell pellet. Cells were washed/resuspended with 40ml milliQ H_2_O, re-pelleted and supernatant was removed. Cells were re-suspended and pooled to a final stock concentration of 6.25×10^8^ cells ml^−1^ based on predictions from previously performed cell extraction and cell counts from the same soils.

#### 2.4.3 Cell counts

2ml cell extract solution was collected for cell quantification at the time of initial blending and after washed cell re-suspension in milliQ H_2_O. Samples were amended gluteraldehyde to give a 1.5% fixation solution and stored at 4ºC for at least 2hrs to allow fixation. Cell counts were carried out using SYBR Green staining and epifluorescence microscopy (Noble and Fuhrman, 1998). Cell solutions were diluted 200 fold, and 6ml was vacuum filtered through 0.2μm Anodisc 25 diameter filters (Whatman, Maidstone, UK). SYBR Green I (Molecular Probes, Eugene, Oregon, Texas) was diluted 2.5×10^−3^ to a working solution. Filters were placed on a 100μL drop of solution and allowed to stain for 20 min in the dark. Filters were oven dried at 60ºC. Duplicate filters per sample were prepared. Filters were mounted onto glass slides with an antifade mounting solution consisting of 50% glycerol, 50% phosphate buffered saline (0.05M Na_2_HPO_4_, 0.85% NaCl, pH 7.5) and 0.1 % p-phenylenediamine. Cells were counted by epifluorescence microscopy.

#### 2.4.4 Inoculation and incubation

All vials used during incubations were placed in a 20ºC waterbath to equilibrate. Headspace overpressure was removed by water filled syringe. All vials were amended with 0.5ml He-flushed NH_4_NO_3_ solution to give a 3mM final concentration. 2ml helium washed concentrated cells from the appropriate soil were added to give a total of ~5×10^7^ cells ml^−1^ in each standard, glutamate amended and carbon negative treatment. 2ml of dummy He flushed milliQ H_2_O was added to make up the volume in cell free chemical extract controls. Vials were magnetically stirred at 360rpm. Headspace gases were sampled and measured every 4hrs using the robotic autosampler gas chromatographs described above under anoxic soil incubations (2.3)

### 2.5 Cell based assay with alternate pH soils

The cell based assay experiment was repeated using soils with contrasting pH and N_2_O hypothetically emitted ratio to test the impact of cells and chemical extract within the context of added pH complexity (Here-after referred to as CBA-pH). Rae’s Junction was used as a high N_2_O hypo emitting low pH (native pH = 5.60, ratio = 0.92) soil, as in CBA-int, while Tapawera was used as the higher pH lower high N_2_O hypothetically emitted (%) soil (native pH 6.58, ratio = 0.68). Again, Rae’s Junction chemical extracts were buffered to pH 6. Tapawera chemical extracts were buffered closer to the native soil pH at 6.6. Triplicate standard treatments and their pHs were: HEC + HEE (6), HEC + LEE (6.6), LEC + HEE (6), LEC + LEE (6.6). Minimum duplicate alternative pH controls were produced for each treatment in which the pH of the treatment chemical extract media was switched to the opposite pH. This allowed determination of the independent effects of pH and chemical extract. Duplicate carbon negative controls and cell negative controls were carried out as in CBA-int but glutamate amended treatments were not included. Full treatment contents and replication is detailed in Table S1.

### 2.6 Nitrate and nitrite quantification

Nitrate + nitrite (NO_3_ ^−^ + NO_2_ ^−^) measurements were performed on soil chemical extracts before incubation media preparation using a previously described chemiluminescent detection method ((Braman and Hendrix, 1989; Lim *et al.*, 2018). This allowed accurate adjustment to a 3mM NO_3_^−^ concentration in the cell based assay media. 10μL of chemical extract was injected into a sealed glass piping system containing heated (95ºC) vanadium chloride solution (50mM VCl_3_, 1M HCl). VCl_3_ reacts rapidly with NO_3_^−^ and NO_2_^−^ at high temperature to produce NO gas. Produced NO is transported via an N_2_ carrier stream to a Sievers Nitric Oxide Analyzer 280i system (GE Analytical Instruments, Boulder, CO, USA). Cell based assay sample NO_2_^−^ concentrations during incubations were quantified using the same chemiluminescence detection system, however, a separate reaction crucible containing NaI (1% w/v NaI in 50% acetic acid, room temperature) was used to specifically target NO_2_^−^. Signal peak areas were calibrated using 10μL injections of a 10-fold KNO_3_ or KNO_2_ dilution series (1mM to 0.001mM). A single rep from each CBA treatment was sampled every ~24hrs (0.15ml) for immediate quantification of accumulated NO ^−^.

### 2.7 N_2_O emission potential

Soil and CBA treatment N_2_O emission potential was evaluated based on two time-integrated measures: N_2_O hypothetically emitted (from here on referred to as N_2_O emitted) and N_2_O hypothetically emitted ratio (from here on referred to as N_2_O ratio). Both measures were developed to account for periods of net N_2_O reconsumption from vial headspace which would not occur in an open system and is therefore not indicative of N_2_O emission potential. N_2_O hypothetically emitted is calculated as the sum of net positive N_2_O accumulations between each sampling point over the course of the incubation + N_2_O lost due to sampling dilution. N_2_O hypothetically emitted ratios are calculated as N_2_O hypothetically emitted/(N_2_O hypothetically emitted + N_2_O emission prevented) where the N_2_O emissions prevented term is the total N_2_ finally accumulated in the vial + loses to sampling and leaks - N_2_ derived from reduction of headspace accumulated N_2_O. This formula can also be applied to soil incubations which include only a single N_2_O accumulation peak and the resulting value is almost equivalent to the N_2_O hypothetically emitted (%) term previously utilized in Highton et al. (2020), differing only in use of cumulative N_2_O (zeroed, sampling dilution and leakage accounted for) in calculations rather than the previously used actual in vial quantities.

Differences in these measures of N_2_O emission potential between treatments were evaluated based on non-overlapping 95% confidence intervals. Independent variable (cell origin, chemical extract origin, pH) effects on N_2_O emitted or ratio were calculated by comparison of relevant treatments and with a specific predicted direction of effect in mind. LE cells, chemical extracts and higher pH (6.6) were expected to decrease N_2_O emitted and ratios while HE cells, chemical extracts and lower pH (6.0) were expected to increase N_2_O emitted and ratios. Expected directions of effect were denoted with a positive value and unexpected with a negative value. When averaged, effects were maintained as positive or negative values unless stated that the effect size given was directionless.

### 2.8 Microbiome composition

DNA was extracted from cell stock and parent soil for each soil to determine extraction bias and community differences between separate cell extracts. For soils, parent soil was collected at the start of the cell extraction protocol and stored at −80ºC until DNA extraction of duplicate 0.25g replicates using the DNeasy powerlyzer powersoil extraction kit (Qiagen, Hilden, Germany). Duplicate 5ml cell stock aliquots were harvested just prior to inoculation of cell based assay treatments and frozen at −80ºC until cell pelleting and DNA extraction.

16S amplicon sequencing of samples was carried out on illumina hiseq using Version 4_13 of the Earth Microbiome Project standard protocol (Caporaso *et al.*, 2012). Sequences are available in the NCBI Sequence Read Archive under the BioProject ID PRJNA678002. Sequence quality control and ASV (Amplicon sequence variant) picking was carried out in R version 3.6.1 (R Core Team, 2016) using the dada2 pipeline version 1.12.1 (Callahan *et al.*, 2016). Taxonomy was assigned using the SILVA database (version 132) (Quast *et al.*, 2013) and the RDP (Ribosomal Database Project) bayesian classifier (Wang *et al.*, 2007).

Sample sequence reads were rarefied 10 times to a depth of 11500 sequences using phyloseq package functions (McMurdie and Holmes, 2013). Independent rarefactions were combined and normalised to the number of rarefactions. Fractional ASV counts were rounded to integers.

#### 2.8.1 Beta diversity and ASV sharing

All beta diversity and ASV sharing plots were generated using ggplot2 version 3.2.1 (Ginestet, 2011) and adjusted with the ggpubr (Kassambara, 2020) and forcats (Wickham, 2020) packages unless otherwise stated. The phyloseq package (McMurdie and Holmes, 2013) was used to calculate and display community composition dissimilarity, the mean number of shared and unique ASVs, and the relative abundance of organisms at the phylum rank with the additional usage of the dplyr (Wickham *et al.*, 2019) and Rmisc (Hope, 2013) packages. Community composition dissimilarity patterns were confirmed using vegan package (Dixon, 2003) ANOSIM and ADONIS tests.

The fold change of ASV abundance differences between extracted cells and soil samples, and its accompanying p-value was generated with the use of the edgeR (Robinson *et al.*, 2009) package to identify significantly changing ASVs with an exact test. P-values were adjusted based on Benjamini-Hochberg p value correction and ASVs were only displayed if their false discover rate (FDR) was below 0.1. ASV Genus taxonomy was only labelled if abundance differed more than 5-fold with a p-value < 1 x 10^−4^.

## Results

### 3.1 Soil and cell based incubations have distinct gas accumulation patterns but relative emission potential is conserved

Denitrification gas (NO, N_2_O, N_2_) kinetics were compared between soil and cell based incubations to determine whether the cell based system accurately modeled the trends observed using soils. Soil incubations (Figure 1A, Figure S1) displayed a single N_2_O accumulation and depletion curve. N_2_O ratios were determined by the sequentiality of N_2_O production and reduction steps as previously described in Highton et al. (2020). In the most extreme cases, close to all added N was accumulated as N_2_O before high rate N_2_ production/N_2_O reduction was initiated, predicting high emissions from an in situ (unsealed) environment.

**Figure 1.**
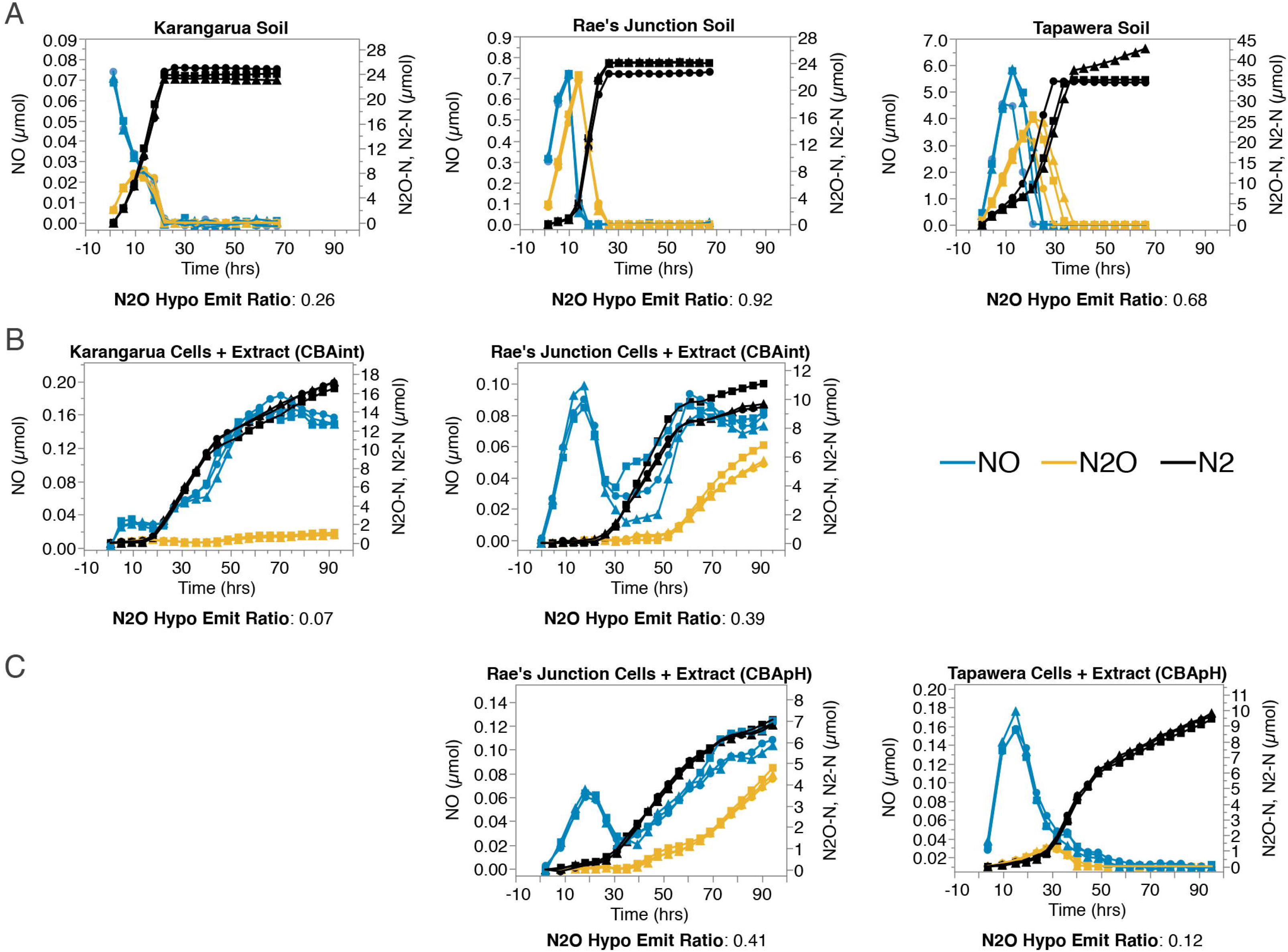
Comparison of parent soils (A) and equivalent unswapped cell based assay treatments (CBA) from CBA-int (B) and CBA-pH (C) reveal contrasting gas accumulation patterns. Headspace gases NO (blue), N_2_O (orange), N_2_ (black) were quantified every 4hrs from triplicate (dots, squares, triangles) 3mM NH_4_NO_3_ amended anoxic incubations. Note separate scales between treatments to highlight relative gas accumulation

Gas accumulation patterns in cell-based incubations were inconsistent with soil incubations. Most treatments experienced an initial lag phase in denitrification product accumulation and CO_2_ accumulation (Figure S2). Only +glutamate treatments completed processing of added N (Figure S3B) during the experimental timeframe. Early N_2_O accumulation was very low while major differences in N_2_O accumulation, and thus N_2_O ratio, occurred later in the incubation when total N turnover rates and N_2_O reduction (N_2_ production) rates suddenly dropped (Figure S3, Figure S4). Late drops in N_2_O reduction rate were usually greater than drops in N_2_O production rates, resulting in increased N_2_O accumulation.

Despite distinct gas accumulation patterns, soil and cell based assays sustained relative rankings based on N_2_O ratios (Figure 2, Rae’s Junct>Tapawera>Karangarua). Gas production profiles were not completely consistent between separate cell based assay runs as evidenced by the repeated Rae’s Junction based incubations (Figure 1, Rae’s Junction vs. 2-Rae’s Junction), however, this variation did not greatly impact N_2_O ratios and relative ranking of incubations (Figure 2).

**Figure 2.**
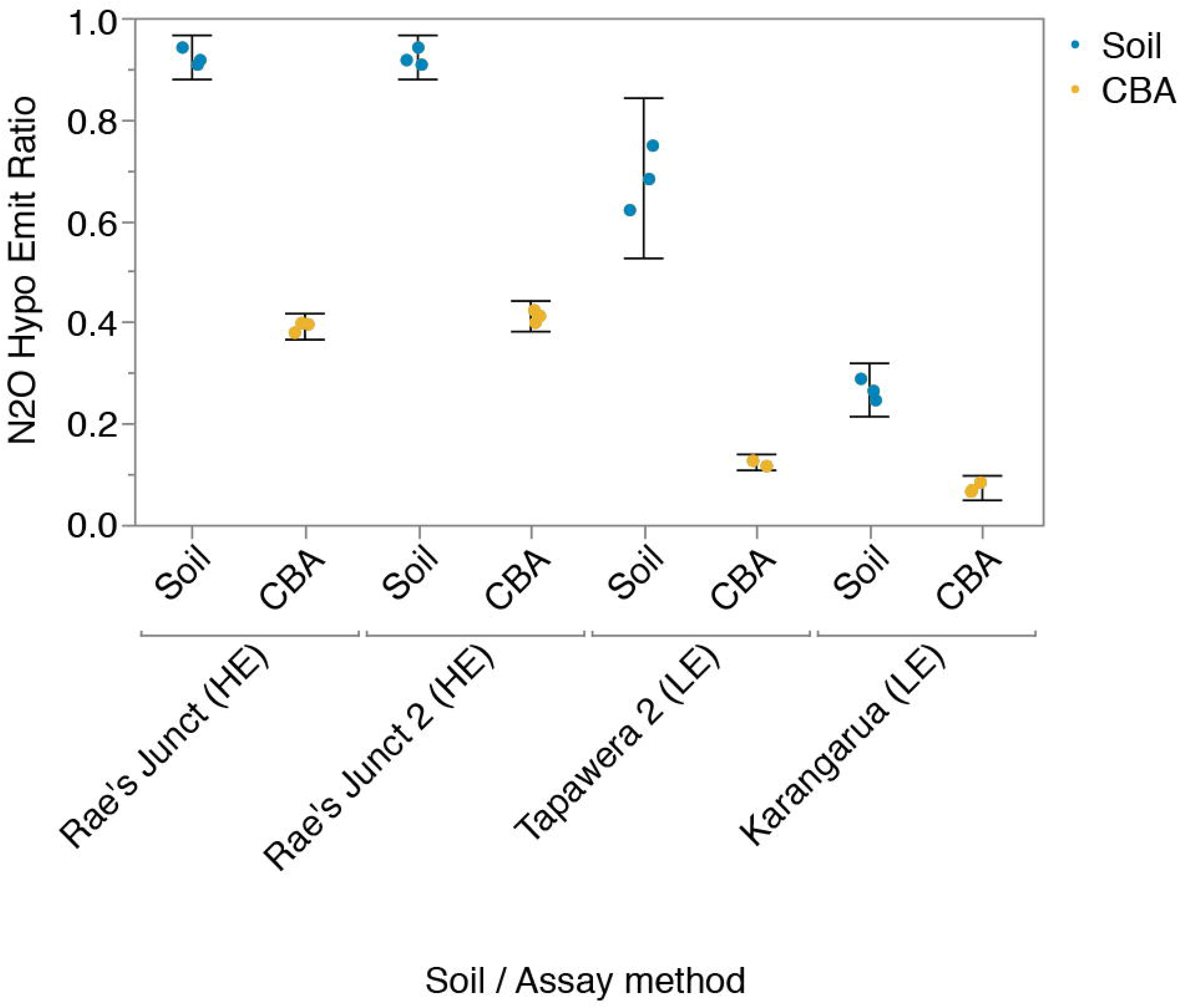
Relative ranking of parent soil N_2_O ratios is maintained in equivalent CBA treatments but lower on an absolute scale. N_2_O ratios summarise the N_2_O emission potential from 90hr CBA anoxic incubations amended with 3mM NH_4_NO_3_ and are calculated as N_2_O/N_2_O+N_2_ at the end of a CBA incubation, where periods of net negative N_2_O accumulation are ignored to account for multiple gas peaks. Equivalent CBA treatments include both cells and chemical extracts derived from the parent soil. Results from triplicate vials per treatment are displayed with 95% confidence intervals.

### 3.2 Both chemistry and microbiome determine N_2_O emission potential

We compared N_2_O ratios and N_2_O accumulation in a CBA (CBA-int) seeded with cells and chemical extracts from soils with similar native pH (5.6, 5.75) to determine whether microbiome (cells) or chemical factors (extracts) were the most important determinant of N_2_O emission potential in the absence of pH effects. Both cell and chemical extract origin affected N_2_O ratio and N_2_O emitted resulting in a gradient: HEC+HEE> LEC+HEE≈HEC+LEE>LEC+LEE (Figure 3A, B). Cell and chemical extract origin had similar impacts on N_2_O ratio but chemical extract origin was the most important determinant of overall emissions, with on average 60% greater impact (Table 1, CBA-int).

**Figure 3.**
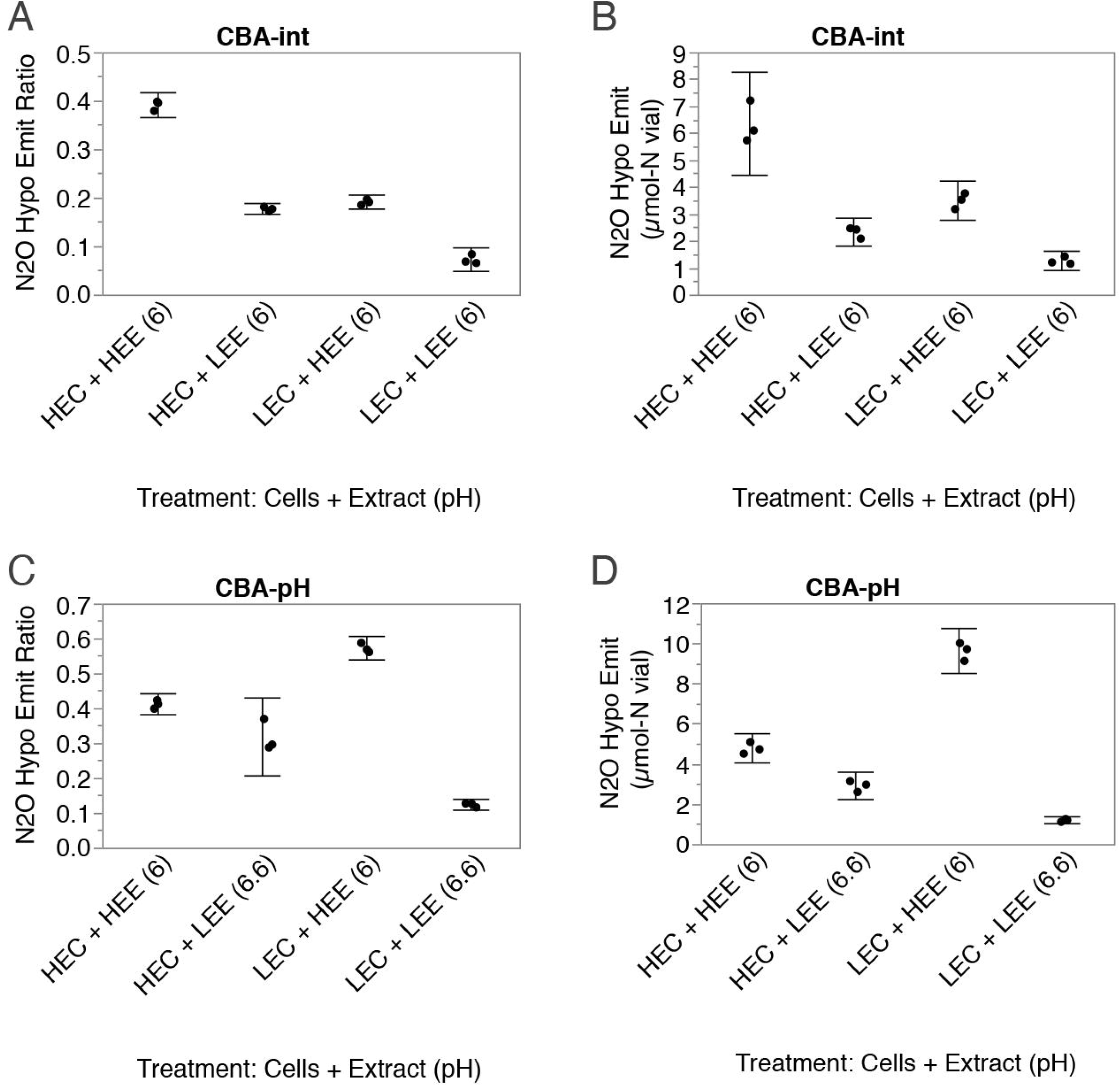
Cell and chemical extract origin impact CBA N_2_O ratios and N_2_O emitted (μmol-N per vial). Standard swap treatments from CBA-int (A,B) or CBA-pH (C,D). N_2_O ratios and N_2_O emitted summarise the N_2_O emission potential from 90hr CBA anoxic incubations amended with 3mM NH_4_NO_3_ and are calculated as N_2_O/N_2_O+N_2_ and total N_2_O accumulated at the end of a CBA incubation, where periods of net negative N_2_O accumulation are ignored to account for multiple gas peaks. Results from triplicate vials per treatment are displayed with 95% confidence intervals. pH of CBA-pH chemical extracts were buffered at two levels and are labeled accordingly.

**Table 1.** Differences in treatment emission potential indicating strength of cell and chemical extract origin effects for CBA-int and CBA-pH

| CBA-int |  |  |  |  | CBA-pH |  |  |  |  |
| --- | --- | --- | --- | --- | --- | --- | --- | --- | --- |
| Treatment | N<sub>2</sub>O hypo<br>emit ratio | 95% CI | N<sub>2</sub>O hypo<br>emit (μmol-N) | 95% CI | Treatment | N<sub>2</sub>O hypo<br>emit ratio | 95% CI | N<sub>2</sub>O hypo<br>emit (μmol-N) | 95% CI |
| HE cells + HE extract | 0.39 | 0.37, 0.42 | 6.34 | 4.43, 8.25 | HE cells + HE extract (6) | 0.41 | 0.38, 0.44 | 4.77 | 4.04, 5.5 |
| HE cells + LE extract | 0.18 | 0.17, 0.19 | 2.32 | 1.8, 2.84 | HE cells + LE extract (6.6) | 0.32 | 0.21, 0.43 | 2.91 | 2.22, 3.58 |
| LE cells + HE extract | 0.19 | 0.18, 0.21 | 3.49 | 2.76, 4.22 | LE cells + HE extract (6) | 0.57 | 0.54, 0.61 | 9.61 | 8.49, 10.73 |
| LE cells + LE extract | 0.07 | 0.05, 0.1 | 1.25 | 0.9, 1.61 | LE cells + LE extract (6.6) | 0.12 | 0.11, 0.14 | 1.19 | 1.01, 1.36 |
| Cell effect | Differences |  | Differences |  | Cell effect | Differences |  | Differences |  |
| HE, HE vs LE, HE | 0.20 | 0.18, 0.22 | 2.85 | 1.19, 4.52 | HE, HE vs LE, HE | -0.16* | -0.19, -0.13 | -4.85* | -5.77, -3.92 |
| HE, LE vs LE, LE | 0.10 | 0.08, 0.12 | 1.07* | 0.64, 1.49 | HE, LE vs LE, LE | 0.19* | 0.09, 0.3 | 1.72* | 1.08, 2.35 |
| Average cell effect | 0.15 |  | 1.96 |  | Average cell effect | 0.02 |  | -1.56 |  |
| Av Directionless cell effect | 0.15 |  | 1.96 |  | Av Directionless cell effect | 0.18 |  | 3.28 |  |
| Extract effect | Differences |  | Differences |  | Extract effect | Differences |  | Differences |  |
| HE, HE vs HE, LE | 0.21 | 0.19, 0.24 | 4.02 | 2.26, 5.78 | HE, HE vs HE, LE | 0.09* | -0.01, 0.2 | 1.86* | 1.22, 2.51 |
| LE, HE vs LE, LE | 0.12 | 0.1, 0.14 | 2.23* | 1.62, 2.84 | LE, HE vs LE, LE | 0.45* | 0.42, 0.48 | 8.43* | 7.34, 9.51 |
| Average extract effect | 0.17 |  | 3.13 |  | Average extract effect | 0.27 |  | 5.14 |  |
| Av Directionless extract effect | 0.17 |  | 3.13 |  | Av Directionless extract effect | 0.27 |  | 5.14 |  |

Emission potential differences are expressed relative to the HE extract or cells. Positive values indicate reduced N<sub>2</sub>O emission potential when comparatively LE extracts or cells were used.
Average cell effects are calculated using signed difference values while average directionless cell effects are calculated using absolute values
\* Indicates difference values have non-overlapping confidence intervals with the appropriate comparison. Direct comparison of cell vs. chemical extract difference values should be compared relative to the equivalent baseline treatment i.e. HE, HE vs. LE, HE compared with HE, HE vs. HE, LE.

To account for the role of pH, soils with differing N_2_O ratio and pH were also compared (CBA-pH). pH of the treatment was coupled to the soil chemical extract (HE extracts: 6.0, LE extracts: 6.6). Again, both cell and chemical extract origin (including coupled pH) affected N_2_O ratio and N_2_O emitted resulting in a gradient: LEC+HEE>HEC+HEE>HEC+LEE>LEC+LEE (Figure 3C, D) but chemical extract was the most important determinant of both N_2_O ratio and emissions (Table 1, CBA-pH).

Patterns were largely determined by the unexpected emission patterns of LE cells which had very high emission potential in the presence of HE extracts yet low emission potential in the presence of LE extracts. **Negative** emission potential difference values (Table 1, CBA-pH) indicate the unexpected **increase** in emission potential using LE cells in the presence of HE extract.

### 3.3 pH has an outsized impact on low emitting cells

pH switched control treatments (HEE 6.0➔6.6, LEE 6.6➔6.0) revealed high N_2_O ratio in the LEC+HEE treatment was largely a response to the low pH of the HE extracts; LE cell N_2_O ratios were much more sensitive to independent pH change than HE cells (Table 2). We accounted for these strong impacts on LE cells by examination of the overall assay at pH 6.6, revealing a similar trend to the CBA-int assay: equal impact of cell and chemical extract origin on ratio (average change of 0.13 points), greater impact of chemical extract on total N_2_O emissions (average change cell= 0.37μmol-N, chemical extract=4.19, Table S2, overall). However, it should be noted that independent impact of HE extracts still lead to unexpectedly high absolute N_2_O emissions from the LE+HE treatment at pH 6.6 due to rate effects of the from the HE extract (Table S2, overall).

**Table 2.** Difference in N2O hypo emit (ratio) associated with independent treatment difference in pH or chemical extract relative to a baseline sample

| Difference in N<sub>2</sub>O hypo emit ratio for treatments varying in: |  |  |  |  |  |  |  |
| --- | --- | --- | --- | --- | --- | --- | --- |
| Baseline sample | Extract | 95% CI | pH | 95% CI | Extract + pH actual | 95% CI | Extract + pH predicted (sum independent extract and pH differences) |
| 6 HEC + HEE | -0.03 | -0.08, 0.01 | -0.03 | -0.33, 0.28 | -0.09 | -0.2, 0.01 | -0.06 |
| 6.6 HEC + LEE | 0.07 | -0.21, 0.08 | 0.06 | -0.03, 0.15 | 0.09 | -0.01, 0.2 | 0.13 |
| 6 LEC + HEE | -0.33 | -0.52, -0.15 | -0.26 | -0.29, -0.23 | -0.45 | -0.48, -0.42 | -0.59 |
| 6.6 LEC + LEE | 0.19 | 0.17, 0.21 | 0.12 | -0.13, 0.36 | 0.45 | 0.42, 0.48 | 0.31 |

| Difference in N<sub>2</sub>O hypo emit (μmol-N) for treatments varying in: |  |  |  |  |  |  |  |
| --- | --- | --- | --- | --- | --- | --- | --- |
| Baseline sample | Extract | 95% CI | pH | 95% CI | Extract + pH actual | 95% CI | Extract + pH predicted (sum independent extract and pH differences) |
| 6 HE + HE | -1.74 | -2.52, -0.96 | 0.98 | -2.31, 4.26 | -1.86 | -2.49, -1.23 | -0.76 |
| 6.6 HE + LE | 2.84 | -0.73, 6.4 | 0.12 | -0.88, 0.65 | 1.86 | 1.23, 2.49 | 2.96 |
| 6 LE + HE | -7.17* | -8.37, -5.99 | -2.88* | -3.85, -1.93 | -8.42 | -9.51, -7.34 | -10.06 |
| 6.6 LE + LE | 5.54* | 5.12, 5.96 | 1.25* | -1.54, 4.04 | 8.42 | 7.34, 9.51 | 6.78 |

Negative values indicate a lower N<sub>2</sub>O ratio or hypothetical emissions relative to the baseline sample
\* Indicates pH and chemical extract difference values have non overlapping confidence intervals for equivalent baseline samples (same row)

Comparison of independent pH, and chemical extract origin effects revealed an additional two notable pH related phenomena:

1. Low pH drove large increases in N_2_O ratio (average change 0.11 points), on par with independent chemical extract effects (Figure 4A), yet only minor changes in total N_2_O emissions (average 1.30 μmol-N, Figure 4B) due to the contrasting impact of pH on N turnover rates and N_2_O ratios. In one instance pH increase to 6.6 actually increased total emissions (Table 2, 6 HEC + HEE).
2. Low pH and HE extract acted synergistically to increase LE cell emission potential i.e. Switching pH and chemical extract of 6.6 LEC + LEE treatment to 6 and HE extracts lead to a greater increase in N_2_O ratio and N_2_O emitted than would be predicted by independent changes in pH or extract alone (Table 2). A much weaker positive synergistic effect (reduction in N_2_O ratio and total N_2_O) of LE extracts and LE pH (higher-6.6) on HE cells was also indicated (Table 2).

**Figure 4.**
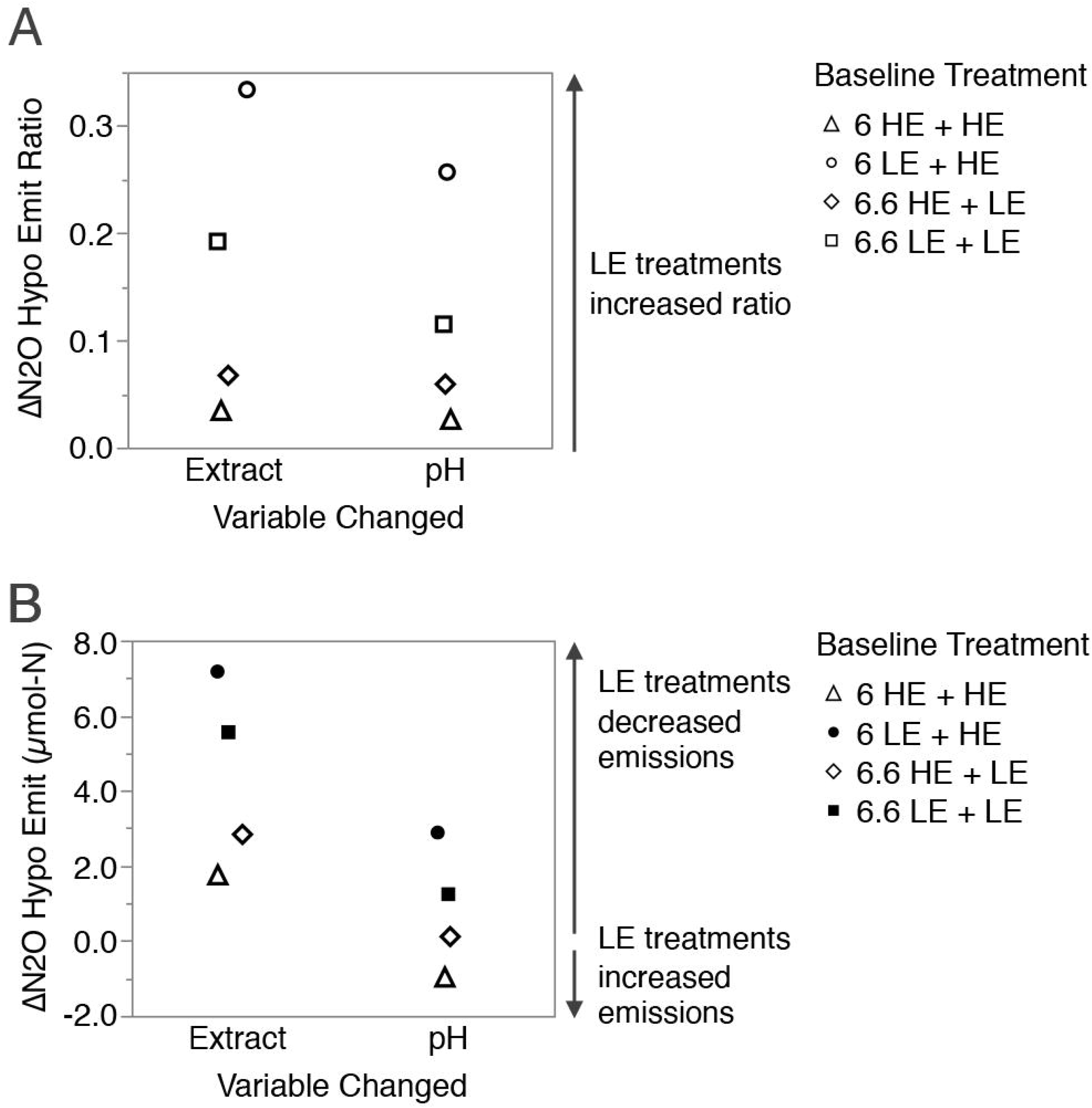
Comparison of independent pH and chemical extract origin changes indicates similar impact of pH and chemical extract on N_2_O ratios (A) but minor impact of pH on N_2_O emitted (B). Each symbol compares the change in N_2_O emission potential from 1 of 4 CBA-pH baseline treatments. Filled symbols indicate non-overlapping 95% confidence intervals for alternative pH or chemical extract changes to the same baseline treatment. Positive values indicate variable change had expected direction of effect on N_2_O ratio or emissions i.e. higher pH and LE extracts are expected to decrease N_2_O ratio and emissions, lower pH and HE extracts vice versa.

### 3.4 Carbon/starvation effect

We hypothesized that sudden changes in N turnover (especially N_2_ production) and emissions during the cell based incubations were linked to shifts in carbon availability. +C (3mM Na-glutamate) controls were included for each swap treatment in CBA-int to determine whether any of the observed differences in treatments were caused by changes in C availability. Divergence of gas accumulation rates in +C controls compared with standard treatments indicated that all treatments became carbon limited during the course of the incubation (Figure 5). Further, carbon amended controls did not experience the late incubation decreases in N_2_ production rate, or the associated increased N_2_O accumulation, seen in -C treatments suggesting these features may result from C limitation. Predicted actual total N gas and CO_2_ production rates typically dropped during the transition to the lower N_2_ rate period also supporting increasing C limitation (Figure S2). CO_2_ rate drops during this time period were often definitive and of high magnitude but were less obvious for some incubations: HEC + LEE, 6 HEC + HEE, 6.6 LEC + LEE. We carried out a further analysis separating the impact of cell and chemical extracts during the carbon non-limited and limited periods of the incubation (Supplemental document S1, Figure S5)

**Figure 5.**
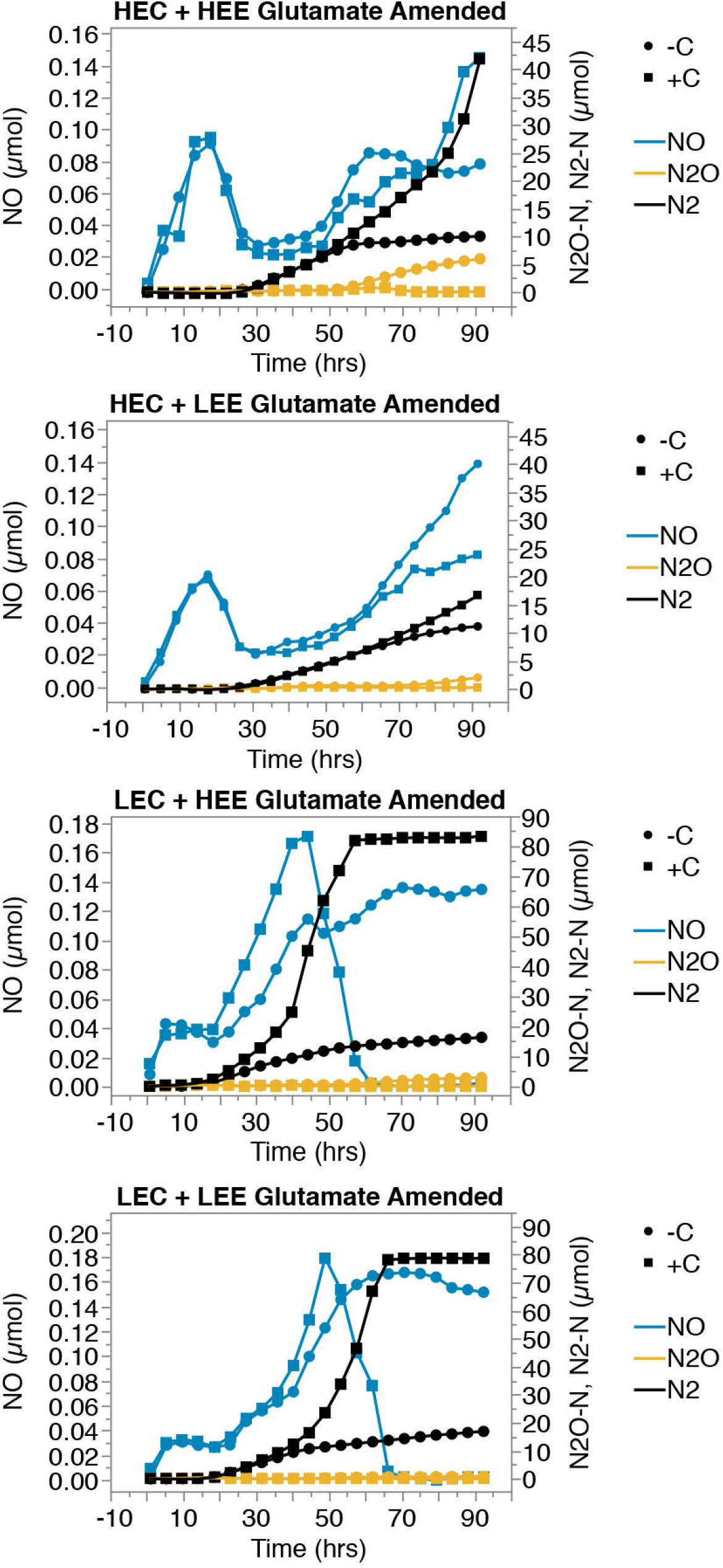
Carbon limitation associated with increased N_2_O accumulation and reduced N_2_ accumulation in CBA-int incubations. Standard treatments (dots), 3mM glutamate amended treatments (squares). Headspace gases NO (blue), N_2_O (orange), N_2_ (black) were quantified every 4hrs from 3mM NH_4_NO_3_ amended anoxic extracted cell and chemistry based incubations. Average gas accumulation from triplicate (standard treatments) or minimum duplicate (glutamate amended treatments) vials per treatment are presented. Note separate scales between treatments to highlight relative gas accumulation.

### 3.5 Microbiome analysis

To assess if extracted cells were representative of soil microbiomes, and to compare differences in microbiomes across soils we used 16S rRNA amplicon sequencing and processed results into amplicon sequence variants (ASVs). Microbiome differences where primarily associated to soil origin (ANOSIM: R^2^◻=◻0.72, p◻<◻0.001, Figure 6A) with extracted cells clustering alongside their original soils. However, small but significant changes were detected between extracted cells and soils (ANOSIM: R^2^◻=◻0.34, p◻=◻0.003). While both extracted cells and soils shared a large proportion (mean 50 % with a standard deviation of 12 %) of their total ASVs (Figure 6B), extracted cells consistently recovered a larger number of ASVs (Wilcox, W = 16, p = 0.029). This bias in ASV detection was reflected at the phylum level (Figure 6C) where Firmicutes where more represented in the soils compared to extract. It also highlighted differences between soils. To identify specific organisms enriched in either soils or extracted cells ASVs with differential abundance between sample type were detected using an exact test (Figure 6D). ASV’s in the Bacillaceae family were significantly enriched in all soils relative to extracted cells but otherwise no consistent extraction bias was observable.

**Figure 6.**
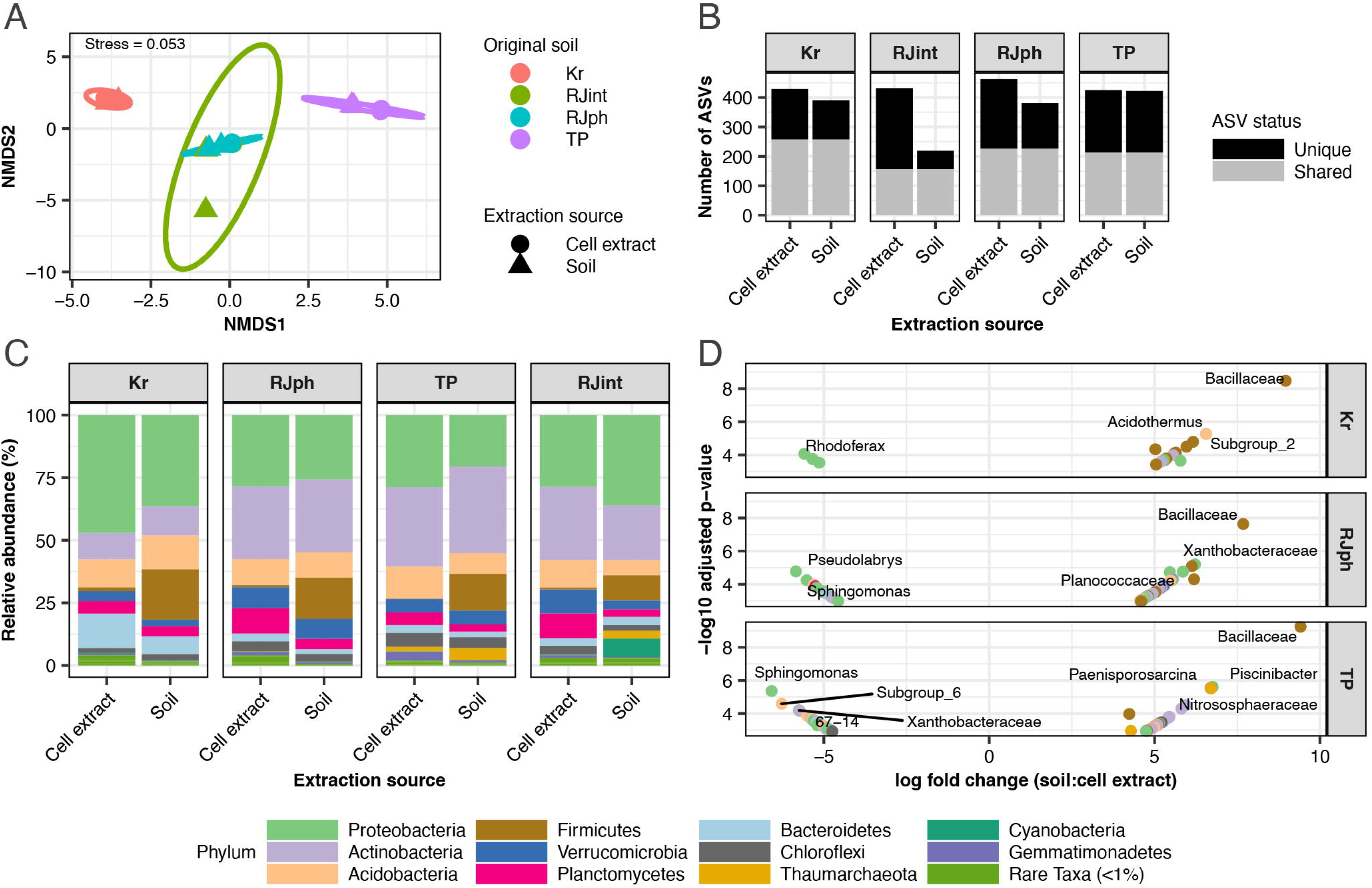
Extraction bias in microbial communities. Community differences due to cell extraction are shown using NMDS (A), zeta-diversity (B), and community abundance (C-D). NMDS shows community dissimilarity (Bray-Curtis), where colours represent origin soil and shapes extraction source (soil or extracted cells). B depicts shared and unique ASVs between soil and cell extracted sequences. C depicts differences in phylum level relative abundance between soil and cell DNA extraction sources. D depicts fold changes in specific ASVs between soil and cell DNA extraction sources, calculated by dividing ASV abundance from soil communities, by those from extracted cells. ASV’s with significant changes are labelled by genera.

## Discussion

### 4.1 Relevance of model to soils

The cell based assay approach allowed causal linkage of microbiome composition and chemistry to N_2_O emission potential. However, as with any model system, applicability to the initial environment studied must be present. Conserved soil rankings based on N_2_O ratios implied general relevance of the system to soils (Figure 2), however, a number of kinetic dissimilarities from soils resulted in different absolute N_2_O ratios, which must be considered.

1. An initial lag phase in which cell based assay incubations accumulated only very low concentrations of CO_2_ and denitrification products NO_2_^−^, NO, N_2_O, N_2_ (Figure S3, Figure S4). This could hypothetically be caused by an initial lack of sufficient denitrifier cell density or a stress response to the cell extraction procedure. Lag or at least very low early denitrification activity and CO_2_ production is also observable in some previous soil-extracted cell based experiments, though the cause is unclear (Nadeem *et al.*, 2013; Brenzinger *et al.*, 2015; Dörsch *et al.*, 2012).
2. Low N_2_O accumulation during the early incubation period (Figure S3, Figure S4). This occurred in most CBA treatments, notably excluding those containing Tapawera cells, and resulted in lowered N_2_O ratios relative to parent soils. Low N_2_O accumulation could be a feature of extracted cell based incubations in the presence of easily utilizable carbon as indicated by very low N_2_O accumulation in the present carbon amended cell based assay treatments (Figure S3B) and a number of previous extracted cell based studies (Dörsch *et al.*, 2012; Brenzinger *et al.*, 2015).
3. A secondary period of high N_2_O accumulation/reduced N_2_ production rates in cell based incubations. Evidence discussed below (4.4) suggests this was most likely a result of carbon limitation and utilization of less energetically favourable carbon sources.

In addition to the explanations given above, the kinetic dissimilarities between soils and cell based incubations are potentially explained by a variety of differences in experimental conditions. Soil and cell incubations most likely differed in cell density and numbers, microbiome composition (due to any biases inherent in the extraction procedure (Holmsgaard *et al.*, 2011; Nadeem *et al.*, 2013)), carbon availability and type (the soluble water extractable component of soil C is usually only around 1% of total soil C and no attempt was made to match carbon concentration in incubations to soils (Gregorich *et al.*, 2003; Guigue *et al.*, 2014)), time of soil in storage (due differences in when separate incubation experiments were carried out), and notably, physical differences including, presence/absence of soil particles, water content and stirring. Water can slow gas diffusion by 4 orders of magnitude (Heincke and Kaupenjohann, 1999), initially leading to gas retention (Clough *et al.*, 2005) while soil heterogeneity might limit or enhance local carbon availability (Parkin, 1987; Kuzyakov and Blagodatskaya, 2015).

Relevance to the soils is further dependent on extracted microbiomes accurately representing soil microbiomes. During any soil cell extraction method, only a portion of soil cells are extracted (Lindahl and Bakken, 1995) leaving the possibility for biases in composition of the community extracted. For example, Nycodenz based extractions have previously been shown to result in reduced microbiome diversity and bias towards or against certain bacterial phyla compared to parent soils (Holmsgaard *et al.*, 2011). Different dispersal methods may also recover metabolically distinct communities (strongly attached vs. loosely attached cells) with different N_2_O emission potentials (Nadeem *et al.*, 2013).

Our own investigations revealed high similarity between parent soil and extracted cell microbiomes at a DNA level (Figure 6A). Unfortunately, we are unable to completely confirm this DNA represented viable cells rather than dead or free floating DNA which passed through the cell extraction procedure. Further, our investigations consistently identified a high number of unique ASVs in extracted cells and total observed richness above that captured from soils. The reason for this is unlikely to be resolved without further empirical evidence but could be due to the larger soil pool and concentration steps used for cell extraction vs. direct soil DNA extractions, movement of species out of rare biosphere in response to the cell extraction protocol disturbance, removal of DNA sorbing soil particles which otherwise inhibit recovery of DNA during extraction (Paulin *et al.*, 2013), dilution of soil pcr/sequencing inhibitors, or increased relative abundance of rarer species due to destruction of abundant organisms during cell extraction. Irrespective of the above limitations, the extracted microbiomes from separate soils will with certainty represent distinct microbiomes from one another, while the conserved relative ranking of N_2_O hypo emit ratios between soil and cell based assays indicate representivity at a functional level (Figure 2).

### 4.2 Proximal vs. microbiome effects

Cell origin impacted both N_2_O ratio and emissions (Table 1), indicating a strong role for microbiome composition in mediating N_2_O emission potential. Previous extracted cell based studies support this claim (Dörsch *et al.*, 2012; Nadeem *et al.*, 2013; Liu *et al.*, 2014) but have typically focused on understanding soil community responses to pH and provide little evaluation of overall impact of community differences compared to other chemical controls. In contrast, another soil based study previously found minimal impact of distal control (implied microbiome composition) on N_2_O ratio but significant impact on total emissions (rate/ enzyme activity) (Čuhel and Šimek, 2011). Here, the directionless effect size of cell origin effects on N_2_O ratio and emissions across both CBAs were not minor, on average only 22 and 37% lower than chemical effects. Therefore, microbiome composition should be considered an important determinant of N_2_O emission potential.

Directional analyses (i.e. LE cells and chemical extracts are expected to decrease N_2_O emission potential and HE cells/extracts vice versa) supported the notion that specific microbiomes and chemical backgrounds can be predictably generalized as lower or higher emitting. In the absence of pH effects (CBA-int or CBA-pH at pH 6.6) LE cells and chemical extracts predictably lowered total emissions and ratios while HE cells and chemical extracts increased them (Table 1, Table S2). Excepting a single case in which HE extracts increased total emissions due to an increased N turnover rate (Table S2, LE cells + HE extract). Such communities or chemical backgrounds might hypothetically be selected for in farms soils to reduce N_2_O emissions. Generalizations might also be applied about the relative importance of microbiome and chemical backgrounds. In the absence of pH effects (CBA-int or CBA-pH at pH 6.6) cell and chemical extracts had a similar average impact on N_2_O ratios but chemical extracts had a greater impact on total emissions due to rate effects (Table 1, Table S2).

Contrastingly, our assays also supported specific less predictable interactions between certain cells, chemical backgrounds and pH that broke the above generalizations. Tapawera LE cells were particularly sensitive to lower pH (Table 2) and especially so in the HE chemical background, showing the highest ratios and total emissions of any treatment (Table 1jhjom CBA-pH). Our ultimate interpretation is that some generalisations can be made about what is a “good” (low N_2_O emitting) denitrifying community and chemical background but that unpredictable specific effects may occur, especially when cells are denitrifying below their typical pH.

An important caveat of all the above interpretations is our inability to completely confirm that cell origin effects were only the result of community composition effects. Extracted cells clearly displayed some lesser but notable activity when incubated in just H_2_O (Figure S3C, Figure S4C) indicating some carbon pool associated with the cells (lysed cells, adherent carbon, stored carbon). Differences in this carbon availability between different cell extractions could potentially influence the denitrification kinetics within the main treatments, especially rates. Cell + H_2_O controls demonstrate similar gas accumulation rates across both cell types in CBA-int (Figure S3C) indicating that, most likely, cell associated carbon should have little observable impact on treatment differences. However, this cannot be claimed for CBA-pH where gas accumulation rates were clearly lower in HE cell + H_2_O controls (Figure S4C).

### 4.3 pH effects

pH differences of just 0.6 points could account for similar changes in N_2_O ratio as differences in chemical extract during CBA-pH (Figure 4A). This is consistent with denitrification literature which commonly identifies pH as a major driving factor of differences in N_2_O/N_2_ emission ratios between soils (Simek and Cooper, 2002; Čuhel and Šimek, 2011; Liu *et al.*, 2014). In contrast, N_2_O emissions were much less susceptible to pH change compared with chemical extract origin due to the conflicting effects of pH on N_2_O ratio and denitrification rates, which are also previously noted (Šimek *et al.*, 2002). In one case, lowering the pH actually resulted in increased N_2_O emissions, therefore, this evidence supports the view that pH manipulation of soil is not necessarily a successful approach to reduce overall N_2_O emissions over a fixed time period. Further, we noted the unideal scenario in which decreasing the pH experienced by higher pH adapted cells had a significant negative impact on N_2_O ratio, while increasing the pH experienced by lower pH adapted cells had only a minor positive impact on N_2_O ratio. In essence, it may be easier for pH change to cause detrimental effects than repair them. Although our pH system may be not be ideal to test this effect. Due to the buffer system used, the low pH soil was already above its natural pH under the low pH treatment.

### 4.4 Differential stages in N_2_O production: the role of carbon

The timing of sudden decreases in CO_2_ production and overall denitrification rates (Figure S2), combined with the lack of late N_2_O accumulation from glutamate amended controls (Figure 5) suggest carbon limitation caused the increased N_2_O accumulation and reduced N_2_ rate observed in the later period of the cell based incubations. If simple carbon limitation was occurring, it is expected that drops in CO_2_ production and denitrification rates would wane gradually over time as carbon concentrations reduced, however, the drops in CO_2_ and denitrification rates were often well defined and rapid. Therefore, we suggest the sudden transitions in rates are the result of exhaustion of a more labile carbon pool and initiation, or maintenance, of consumption of a more recalcitrant carbon pool. Soil extracted carbon is typically quantified in these two separate pools with separate consumption rate constants assigned to the consumption of each pool e.g. (Bowen *et al.*, 2009; Guigue *et al.*, 2014; Kalbitz *et al.*, 2003). The multiple (greater than two) N_2_ rate switches observable in some incubations (Figure S3A, LEC + HEE, Figure S6, 6 LEC + LEE extended) suggest effects to denitrification rates could be through greater than two distinct carbon pools of consecutively reduced energy availability.

Alternatively, denitrification rates may be sustained by consumption of energy storage molecules during the reduced N_2_ rate period. Increased N_2_O accumulation was previously shown in monocultures of *Alcaligenes faecalis* during carbon limitation and co-occurred with consumption of energy storage molecules (Schalk-Otte *et al.*, 2000). This was attributed to competition for limited electrons between N_2_O reductase and the previous denitrification reductases. Under this mechanism, differing N-reductase electron carrier affinities or regulatory mechanisms create an uneven distribution of electrons to the separate denitrification steps (Pan *et al.*, 2013; Ribera-Guardia *et al.*, 2014; Wang *et al.*, 2018; Schalk-Otte *et al.*, 2000). Earlier N-reductases are thought to outcompete N_2_O reductase resulting in N_2_O accumulation during limited electron supply. Electron supply can be limited due to substrate availability but also carbon oxidation rates (Pan *et al.*, 2013) which depend on the substrate being utilized (Ribera-Guardia *et al.*, 2014) and presumably the organism carrying out the oxidation.

Electron competition is consistent with concurrent drops in CO_2_ production, N turnover rates and uneven rebalancing of N_2_O production/reduction in the present study, whether this is during consumption of energy storage molecules or more recalcitrant carbon. However, it is unclear how this mechanism should proceed in a complex community of denitrifiers as competition for electrons is only hypothetically viable when N_2_O production and reduction proceed within the same organism. This is not necessarily a valid assumption in a complex denitrifying community where multiple species of denitrifiers could specialize in separate steps of the process due to the modularity of denitrification genes (Graf *et al.*, 2014; Roco *et al.*, 2017; Lycus *et al.*, 2017). Electron competition between N-reductases has been tested in complex communities (Pan *et al.*, 2013; Ribera-Guardia *et al.*, 2014; Wang *et al.*, 2018) and in some cases it was assumed that denitrification was carried out by complete denitrifiers based on the genera of the dominant microbes within the culture (Pan *et al.*, 2013; Wang *et al.*, 2018). In depth sequencing of metagenomes and metatranscriptomes with genome reconstruction would be necessary to actually resolve the modularity of active denitrifiers within the present system since phylogeny is usually considered a poor predictor of denitrification genetic potential (Jones *et al.*, 2008).

A point of confusion, possibly contradicting the above interpretations, is that cell + H_2_O treatments also demonstrated the distinct denitrification rate changes which we have attributed to carbon limitation (Figure S3C, Figure S4C). This either means the carbon limitation hypothesis and associated interpretations are wrong or that these incubations begun with a non or initially less limiting availability of carbon. Cells were washed multiple times during extraction to remove carbon from the suspension solution. It is therefore most likely that the utilized carbon sources in these treatments is derived from lysed cellular constituents, cell adherent carbon, insoluble carbon or stored carbon.

### 4.5 Conclusion

These investigations provide causal evidence for microbiome composition effects on N_2_O emission potential, but these were on average still weaker than chemical effects. Differences in cell based assay gas accumulation kinetics reduce the general applicability of this system to soils but also serendipitously provide evidence that carbon limitation or switching to more recalcitrant carbon sources can lead to increased N_2_O emissions. Investigations into the effects of pH corroborate the large body of research suggesting that this is a particularly important determinant of soil N_2_O emission ratios but also suggest that its impact on total N_2_O emissions over a fixed time period could be minor compared to other soil variables. Ultimately, we add to the mounting evidence that microbiome composition needs to be considered during soil manipulations aimed at reducing N_2_O emissions.

