## Supplementary tables and figures for "Soil Chemistry and Microbiome Determine N_2_O Emission Potential in Soils"

### Supplementary Tables, Figures and Documents

**Table S1.** Cell based assay treatment constituen**ts**

| CBA#1: Parent soils Rae's Junction (RJ) Karangarua (Kr) | | | | | |
|---|---|---|---|---|---|
| **Treatment (pH, cells + extract)** | **$NH_4NO_3$ (final 3 mM conc)** | **Buffered carbon extract** | **pH** | **Replicates** | **Cells** |
| **Standard treatments** | | | | | |
| HEC + HEE | 0.5 ml | 22.5 ml RJ | 6 | 3 | 2 ml RJ |
| HEC + LEE | 0.5 ml | 22.5 ml Kr | 6 | 3 | 2 ml RJ |
| LEC + HEE | 0.5 ml | 22.5 ml RJ | 6 | 3 | 2 ml Kr |
| LEC + LEE | 0.5 ml | 22.5 ml Kr | 6 | 3 | 2 ml Kr |
| **Carbon amended controls** | | | | | |
| HEC + HEE +C | 0.5 ml | 22.5 ml RJ + glutamate (3 mM) | 6 | 2 | 2 ml RJ |
| HEC + LEE +C | 0.5 ml | 22.5 ml Kr + glutamate (3 mM) | 6 | 3 | 2 ml RJ |
| LEC + HEE +C | 0.5 ml | 22.5 ml RJ + glutamate (3 mM) | 6 | 3 | 2 ml Kr |
| LEC + LEE +C | 0.5 ml | 22.5 ml Kr + glutamate (3 mM) | 6 | 2 | 2 ml Kr |
| **Carbon negative controls** | | | | | |
| HEC + $H_2O$ | 0.5 ml | 22.5 ml buffered milliQ $H_2O$ | 6 | 2 | 2 ml RJ |
| LEC + $H_2O$ | 0.5 ml | 22.5 ml buffered milliQ $H_2O$ | 6 | 2 | 2 ml Kr |
| **Cell free controls** | | | | | |
| H extract | 0.5 ml | 22.5 ml RJ | 6 | 2 | 2 ml milliQ $H_2O$ |
| L extract | 0.5 ml | 22.5 ml Kr | 6 | 2 | 2 ml milliQ $H_2O$ |

| CBA#2: Parent soils Rae's Junction (RJ) and Tapawera (Tp) | | | | | |
|---|---|---|---|---|---|
| **Treatment (pH, cells + extract)** | **$NH_4NO_3$ (final 3 mM conc)** | **Buffered carbon extract** | **pH** | **Replicates** | **Cells** |
| **Standard treatments** | | | | | |
| 6 HEC + HEE | 0.5 ml | 22.5 ml RJ | 6 | 3 | 2 ml RJ |
| 6.6 HEC + LEE | 0.5 ml | 22.5 ml Tp | 6.6 | 3 | 2 ml RJ |
| 6 LEC + HEE | 0.5 ml | 22.5 ml RJ | 6 | 3 | 2 ml Tp |
| 6.6 LEC + LEE | 0.5 ml | 22.5 ml Tp | 6.6 | 3 | 2 ml Tp |
| **Alternative pH controls** | | | | | |
| 6.6 HEC + HEE | 0.5 ml | 22.5 ml RJ | 6.6 | 2 | 2 ml RJ |
| 6 HEC + LEE | 0.5 ml | 22.5 ml Tp | 6 | 3 | 2 ml RJ |
| 6.6 LEC + HEE | 0.5 ml | 22.5 ml RJ | 6.6 | 3 | 2 ml Tp |
| 6 LEC + LEE | 0.5 ml | 22.5 ml Tp | 6 | 2 | 2 ml Tp |
| **Carbon negative controls** | | | | | |
| 6 HEC + $H_2O$ | 0.5 ml | 22.5 ml buffered milliQ $H_2O$ | 6 | 2 | 2 ml RJ |
| 6.6 LEC + $H_2O$ | 0.5 ml | 22.5 ml buffered milliQ $H_2O$ | 6.6 | 2 | 2 ml Tp |
| **Cell free controls** | | | | | |
| H extract | 0.5 ml | 22.5 ml RJ | 6 | 2 | 2 ml milliQ $H_2O$ |
| L extract | 0.5 ml | 22.5 ml Tp | 6.6 | 2 | 2 ml milliQ $H_2O$ |

**Table S2.** CBA-pH high pH (6.6) treatments: Differences in treatment emission potential indicating strength of cell and chemical extract origin effects

| | Overall | | | | Period 1 | | | | Period 2 | | | |
|---|---|---|---|---|---|---|---|---|---|---|---|---|
| **Treatment** | **$N_2O$ hypo emit ratio** | **95% CI** | **$N_2O$ hypo emit (µmol-N)** | **95% CI** | **$N_2O$ hypo emit ratio** | **95% CI** | **$N_2O$ hypo emit (µmol-N)** | **95% CI** | **$N_2O$ hypo emit ratio** | **95% CI** | **$N_2O$ hypo emit (µmol-N)** | **95% CI** |
| HE cells + HE extract | 0.38 | 0.01, 0.76 | 5.75 | 0.38, 11.11 | 0.05 | 0, 0.1 | 0.31 | -0.04, 0.66 | 0.65 | -0.03, 1.33 | 5.44 | 0.43, 10.45 |
| HE cells + LE extract | 0.32 | 0.21, 0.43 | 2.91 | 2.27, 3.55 | 0.06 | -0.09, 0.22 | 0.31 | -0.59, 1.21 | 0.53 | 0.34, 0.71 | 2.60 | 1.68, 3.52 |
| LE cells + HE extract | 0.32 | 0.29, 0.34 | 6.73 | 6.25, 7.21 | 0.20 | 0.17, 0.23 | 2.11 | 1.9, 2.33 | 0.43 | 0.38, 0.47 | 4.61 | 3.98, 5.25 |
| LE cells + LE extract | 0.12 | 0.11, 0.14 | 1.19 | 1.01, 1.36 | 0.19 | 0.16, 0.22 | 1.19 | 1.02, 1.36 | 0.00 | 0, 0 | 0.00 | 0, 0 |
| **Cell effect** | **Differences** | | **Differences** | | **Differences** | | **Differences** | | **Differences** | | **Differences** | |
| HE, HE vs LE, HE | 0.07 | -0.08, 0.21 | -0.98 | -5.14, 3.18 | -0.15 | -0.18, -0.13 | -1.80 | -1.99, -1.62 | 0.22 | -0.37, 0.81 | 0.82 | -2.4, 4.05 |
| HE, LE vs LE, LE | 0.19 | 0.09, 0.3 | 1.72 | 1.13, 2.31 | -0.13 | -0.28, 0.02 | -0.87 | -1.73, -0.02 | 0.53 | 0.34, 0.71 | 2.60 | 1.67, 3.52 |
| Average cell effect | 0.13 | | 0.37 | | -0.14 | | -1.34 | | 0.37 | | 1.71 | |
| **Extract effect** | **Differences** | | **Differences** | | **Differences** | | **Differences** | | **Differences** | | **Differences** | |
| HE, HE vs HE, LE | 0.07 | -0.25, 0.38 | 2.84 | -0.73, 6.4 | -0.02 | -0.17, 0.14 | 0.00 | -0.88, 0.88 | 0.12 | -0.15, 0.38 | 2.84 | 0.38, 5.3 |
| LE, HE vs LE, LE | 0.19 | 0.17, 0.21 | 5.54 | 5.12, 5.96 | 0.01 | -0.02, 0.03 | 0.93 | 0.75, 1.11 | 0.43 | 0.38, 0.47 | 4.61 | 3.98, 5.25 |
| Average extract effect | 0.13 | | 4.19 | | 0.00 | | 0.46 | | 0.27 | | 3.73 | |

Emission potential differences are expressed relative to the HE extract or cells. Positive values indicate reduced $N_2O$ emission potential when LE extracts or cells were used

Greyed difference values have non overlapping confidence intervals with the appropriate comparison. Direct comparison of cell vs chemical extract difference values should be compared relative to the equivalent baseline treatment i.e. HE, HE vs. LE, HE compared with HE, HE vs. HE, LE.

**Table S3.** CBA-pH high pH (6) treatments: Differences in treatment emission potential indicating strength of cell and chemical extract origin effects

| | Overall | | | | Period 1 | | | | Period 2 | | | |
|---|---|---|---|---|---|---|---|---|---|---|---|---|
| **Treatment** | **$N_2O$ hypo emit ratio** | **95% CI** | **$N_2O$ hypo emit (µmol-N)** | **95% CI** | **$N_2O$ hypo emit ratio** | **95% CI** | **$N_2O$ hypo emit (µmol-N)** | **95% CI** | **$N_2O$ hypo emit ratio** | **95% CI** | **$N_2O$ hypo emit (µmol-N)** | **95% CI** |
| HE cells + HE extract | 0.41 | 0.38, 0.44 | 4.77 | 4.04, 5.5 | 0.20 | 0.15, 0.26 | 1.27 | 0.93, 1.62 | 0.65 | 0.63, 0.67 | 3.49 | 3.06, 3.93 |
| HE cells + LE extract | 0.38 | 0.32, 0.43 | 3.03 | 2.1, 3.96 | 0.11 | 0.08, 0.14 | 0.48 | 0.34, 0.62 | 0.71 | 0.59, 0.84 | 2.55 | 1.51, 3.58 |
| LE cells + HE extract | 0.57 | 0.54, 0.61 | 9.61 | 8.49, 10.73 | 0.46 | 0.4, 0.51 | 4.62 | 3.95, 5.3 | 0.75 | 0.74, 0.76 | 4.99 | 4.54, 5.44 |
| LE cells + LE extract | 0.24 | -0.04, 0.52 | 2.44 | -0.67, 5.54 | 0.24 | 0.01, 0.48 | 2.44 | -0.67, 5.54 | 0.00 | 0, 0 | 0.00 | 0, 0 |
| **Cell effect** | **Differences** | | **Differences** | | **Differences** | | **Differences** | | **Differences** | | **Differences** | |
| HE, HE vs LE, HE | -0.16 | -0.19, -0.13 | -4.85 | -5.77, -3.92 | -0.25 | -0.3, -0.2 | -3.35 | -3.91, -2.78 | -0.10 | -0.11, -0.08 | -1.50 | -1.9, -1.1 |
| HE, LE vs LE, LE | 0.14 | 0.01, 0.27 | 0.59 | -0.6, 1.78 | -0.13 | -0.27, 0.01 | -1.95 | -4.84, 0.94 | 0.71 | 0.59, 0.84 | 2.55 | 1.51, 3.58 |
| Average cell effect | -0.01 | | -2.13 | | -0.19 | | -2.65 | | 0.31 | | 0.52 | |
| **Extract effect** | **Differences** | | **Differences** | | **Differences** | | **Differences** | | **Differences** | | **Differences** | |
| HE, HE vs HE, LE | 0.03 | -0.01, 0.08 | 1.74 | 0.96, 2.52 | 0.10 | 0.05, 0.14 | 0.79 | 0.49, 1.09 | -0.06 | -0.19, 0.06 | 0.95 | 0.06, 1.84 |
| LE, HE vs LE, LE | 0.33 | 0.15, 0.52 | 7.18 | 5.99, 8.37 | 0.21 | 0.12, 0.31 | 2.19 | 0.83, 3.55 | 0.75 | 0.74, 0.76 | 4.99 | 4.54, 5.44 |
| Average extract effect | 0.18 | | 4.46 | | 0.16 | | 1.49 | | 0.34 | | 2.97 | |

Emission potential differences are expressed relative to the HE extract or cells. Positive values indicate reduced $N_2O$ emission potential when LE extracts or cells were used

Greyed difference values have non overlapping confidence intervals with the appropriate comparison. Direct comparison of cell vs chemical extract difference values should be compared relative to the equivalent baseline treatment i.e. HE, HE vs. LE, HE compared with HE, HE vs. HE, LE.

**Table S4.** CBA-int: Differences in treatment emission potential indicating strength of cell and chemical extract origin effects

| | **Overall** | | | | **Period 1** | | | | **Period 2** | | | |
|---|---|---|---|---|---|---|---|---|---|---|---|---|
| **Treatment** | **$N_2O$ hypo emit ratio** | **95% CI** | **$N_2O$ hypo emit (μmol-N)** | **95% CI** | **$N_2O$ hypo emit ratio** | **95% CI** | **$N_2O$ hypo emit (μmol-N)** | **95% CI** | **$N_2O$ hypo emit ratio** | **95% CI** | **$N_2O$ hypo emit (μmol-N)** | **95% CI** |
| HE cells + HE extract | 0.39 | 0.37, 0.42 | 6.34 | 4.43, 8.25 | 0.12 | 0, 0.22 | 1.03 | 0.02, 2.03 | 0.75 | 0.71, 0.79 | 5.31 | 2.42, 8.21 |
| HE cells + LE extract | 0.18 | 0.17, 0.19 | 2.32 | 1.8, 2.84 | 0.11 | 0.03, 0.19 | 1.17 | 0.35, 2 | 0.51 | 0.3, 0.72 | 1.15 | 0.05, 2.25 |
| LE cells + HE extract | 0.19 | 0.18, 0.21 | 3.49 | 2.76, 4.22 | 0.09 | 0.08, 0.08 | 0.93 | 0.72, 1.15 | 0.39 | 0.26, 0.5 | 2.56 | 2.02, 3.09 |
| LE cells + LE extract | 0.07 | 0.05, 0.1 | 1.25 | 0.9, 1.61 | 0.04 | 0.03, 0.04 | 0.38 | 0.28, 0.47 | 0.12 | 0.09, 0.14 | 0.88 | 0.43, 1.32 |
| **Cell effect** | **Differences** | | **Differences** | | **Differences** | | **Differences** | | **Differences** | | **Differences** | |
| HE, HE vs LE, HE | 0.20 | 0.18, 0.22 | 2.85 | 1.19, 4.52 | 0.03 | -0.08, 0.14 | 0.10 | -0.85, 1.04 | 0.35 | 0.25, 0.47 | 2.76 | -0.01, 5.53 |
| HE, LE vs LE, LE | 0.10 | 0.08, 0.12 | 1.07 | 0.64, 1.49 | 0.07 | -0.01, 0.15 | 0.80 | -0.01, 1.61 | 0.39 | 0.18, 0.6 | 0.27 | -0.68, 1.22 |
| Average cell effect | 0.15 | | 1.96 | | 0.05 | | 0.45 | | 0.37 | | 1.51 | |
| **Extract effect** | **Differences** | | **Differences** | | **Differences** | | **Differences** | | **Differences** | | **Differences** | |
| HE, HE vs HE, LE | 0.21 | 0.19, 0.24 | 4.02 | 2.26, 5.78 | 0.00 | -0.09, 0.1 | -0.15 | -1, 0.7 | 0.24 | 0.03, 0.44 | 4.17 | 1.64, 6.69 |
| LE, HE vs LE, LE | 0.12 | 0.1, 0.14 | 2.23 | 1.62, 2.84 | 0.05 | 0.04, 0.05 | 0.56 | 0.37, 0.74 | 0.27 | 0.15, 0.38 | 1.68 | 1.22, 2.13 |
| Average extract effect | 0.17 | | 3.13 | | 0.02 | | 0.21 | | 0.26 | | 2.92 | |

Emission potential differences are expressed relative to the HE extract or cells. Positive values indicate reduced $N_2O$ emission potential when LE extracts or cells were used

Greyed difference values have non overlapping confidence intervals with the appropriate comparison. Direct comparison of cell vs chemical extract difference values should be compared relative to the equivalent baseline treatment i.e. HE, HE vs. LE, HE compared with HE, HE vs. HE, LE.

**Table S5.** CBA-pH: Differences in treatment emission potential indicating strength of cell and chemical extract origin effects

| | Overall | | | | Period 1 | | | | Period 2 | | | |
|---|---|---|---|---|---|---|---|---|---|---|---|---|
| **Treatment** | **$N_2O$ hypo emit ratio** | **95% CI** | **$N_2O$ hypo emit (µmol-N)** | **95% CI** | **$N_2O$ hypo emit ratio** | **95% CI** | **$N_2O$ hypo emit (µmol-N)** | **95% CI** | **$N_2O$ hypo emit ratio** | **95% CI** | **$N_2O$ hypo emit (µmol-N)** | **95% CI** |
| 6 HE cells + HE extract | 0.41 | 0.38, 0.44 | 4.77 | 4.04, 5.5 | 0.20 | 0.16, 0.25 | 1.27 | 0.93, 1.62 | 0.65 | 0.64, 0.67 | 3.49 | 3.06, 3.93 |
| 6.6 HE cells + LE extract | 0.32 | 0.21, 0.43 | 2.91 | 2.22, 3.58 | 0.06 | -0.09, 0.21 | 0.31 | -0.59, 1.21 | 0.53 | 0.34, 0.71 | 2.60 | 1.68, 3.52 |
| 6 LE cells + HE extract | 0.57 | 0.54, 0.61 | 9.61 | 8.49, 10.73 | 0.46 | 0.4, 0.51 | 4.62 | 3.95, 5.3 | 0.75 | 0.74, 0.76 | 4.99 | 4.55, 5.44 |
| 6.6 LE cells + LE extract | 0.12 | 0.11, 0.14 | 1.19 | 1.01, 1.36 | 0.19 | 0.16, 0.22 | 1.19 | 1.01, 1.36 | 0.00 | 0, 0 | 0.00 | 0, 0 |
| **Cell effect** | **Differences** | | **Differences** | | **Differences** | | **Differences** | | **Differences** | | **Differences** | |
| HE, HE vs LE, HE | -0.16 | -0.19, -0.13 | -4.85 | -5.77, -3.92 | -0.25 | -0.3, -0.2 | -3.35 | -3.91, -2.78 | -0.10 | -0.11, -0.08 | -1.50 | -1.9, -1.1 |
| HE, LE vs LE, LE | 0.19 | 0.09, 0.3 | 1.72 | 1.08, 2.35 | -0.13 | -0.27, 0.01 | -0.87 | -1.73, -0.02 | 0.53 | 0.34, 0.71 | 2.60 | 1.67, 3.52 |
| Average cell effect | 0.02 | | -1.56 | | -0.19 | | -2.11 | | 0.21 | | 0.55 | |
| **Extract effect** | **Differences** | | **Differences** | | **Differences** | | **Differences** | | **Differences** | | **Differences** | |
| HE, HE vs HE, LE | 0.09 | -0.01, 0.2 | 1.86 | 1.22, 2.51 | 0.14 | 0.01, 0.28 | 0.96 | 0.18, 1.74 | 0.13 | -0.06, 0.31 | 0.90 | 0.12, 1.67 |
| LE, HE vs LE, LE | 0.45 | 0.42, 0.48 | 8.43 | 7.34, 9.51 | 0.26 | 0.22, 0.31 | 3.44 | 2.81, 4.07 | 0.75 | 0.74, 0.76 | 4.99 | 4.54, 5.44 |
| Average extract effect | 0.27 | | 5.14 | | 0.20 | | 2.20 | | 0.44 | | 2.94 | |

Emission potential differences are expressed relative to the HE extract or cells. Positive values indicate reduced $N_2O$ emission potential when LE extracts or cells were used

Greyed difference values have non overlapping confidence intervals with the appropriate comparison. Direct comparison of cell vs chemical extract difference values should be compared relative to the equivalent baseline treatment i.e. HE, HE vs. LE, HE compared with HE, HE vs. HE, LE.

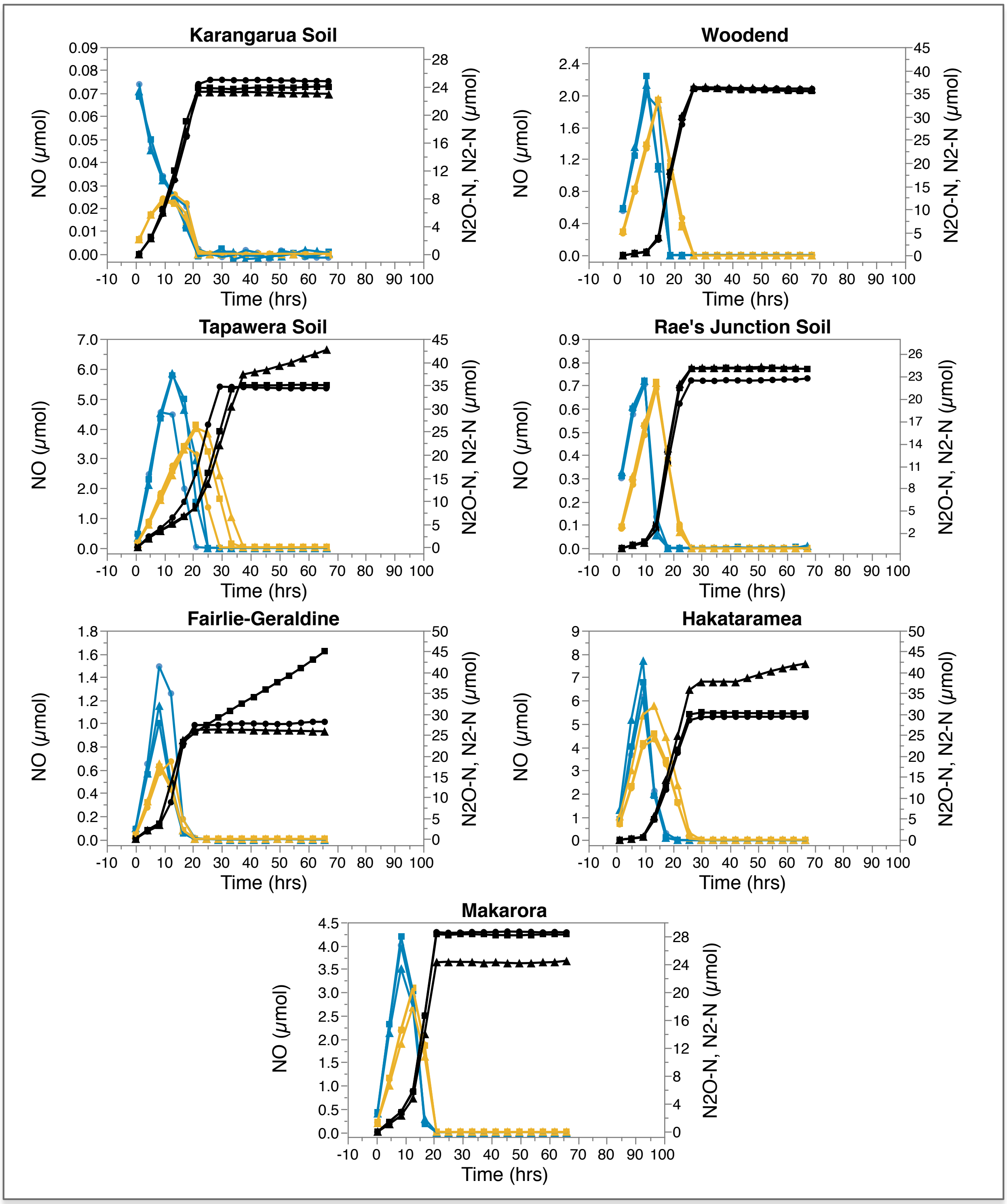


**Figure S1.** Soil denitrification gas accumulation kinetics show contrasting $N_2O$ emission potential based on early $N_2O$ reduction ($N_2$ production) activity. Soils are ordered anticlockwise from low to higher $N_2O$ hypo emit ratios. Headspace gases NO (blue), $N_2O$ (orange), $N_2$ (black) were quantified every 4hrs from triplicate (dots, squares, triangles) 3mM $NH_4NO_3$ amended (by flooding and draining) anoxic soil incubations. Note separate scales between treatments to highlight relative gas accumulation.

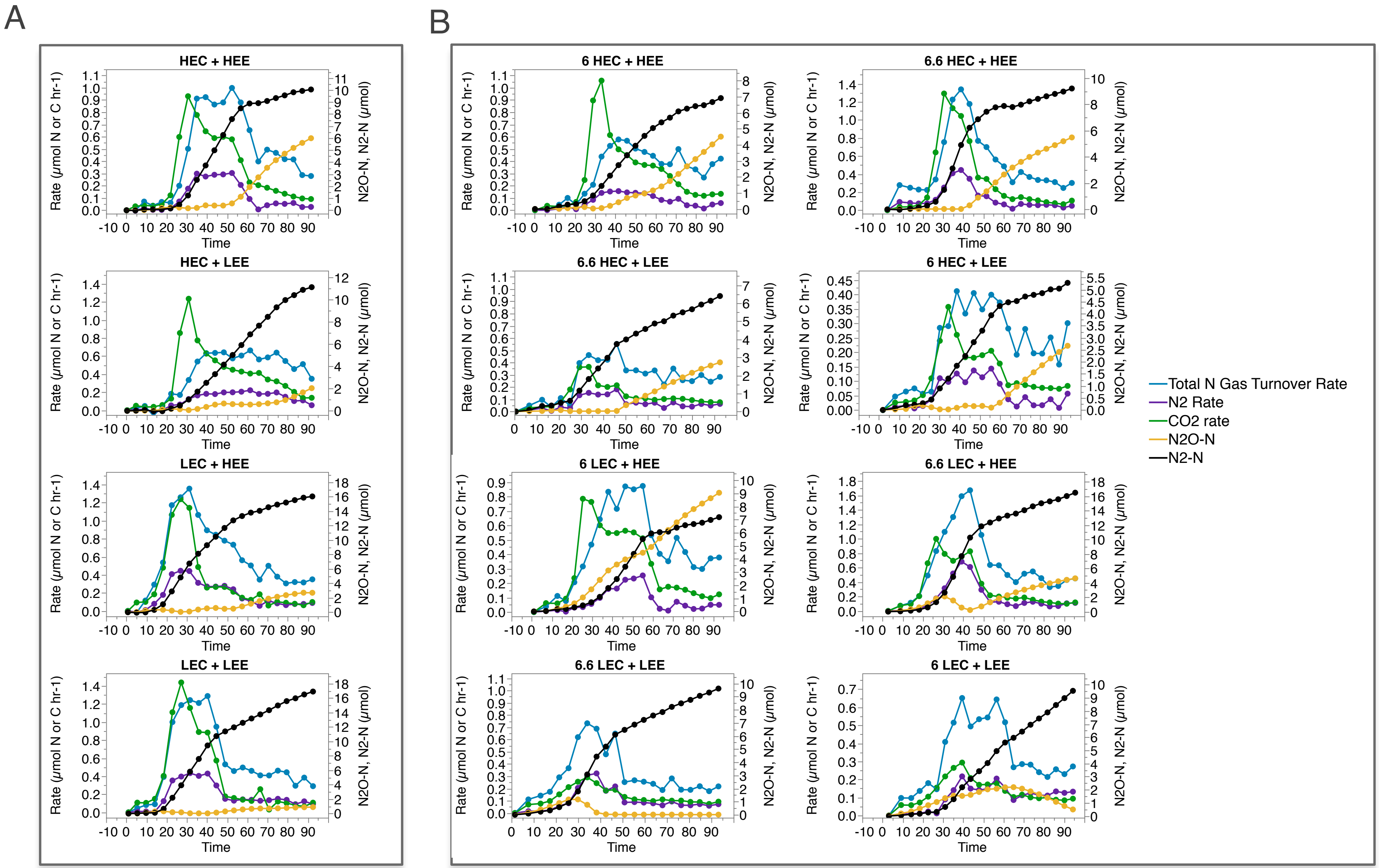
A
B
HEC + HEE
HEC + LEE
LEC + HEE
LEC + LEE
6 HEC + HEE
6.6 HEC + HEE
6.6 HEC + LEE
6 HEC + LEE
6 LEC + HEE
6.6 LEC + HEE
6.6 LEC + LEE
6 LEC + LEE
Rate (μmol N or C hr-1)
N2O-N, N2-N (μmol)
Time
Total N Gas Turnover Rate
N2 Rate
CO2 rate
N2O-N
N2-N

**Figure S2.** Sudden reductions in N gas turnover rate and $CO_2$ accumulation rates are associated with sudden decreases in $N_2$ accumulation rate and increased $N_2O$ accumulation. CBA-int standard treatments (A) and CBA-pH standard treatments + pH controls (B). Headspace gases NO (blue), $N_2O$ (orange), $N_2$ (black) were quantified every 4hrs from 3mM $NH_4NO_3$ amended anoxic extracted cell and carbon based incubations. Average gas accumulation and rates from triplicate or minimum duplicate (alternative pH controls, B right) incubations are presented for total N gas turnover rate (blue), $N_2$ accumulation rate (purple), $CO_2$ accumulation rate (green), $N_2O$ accumulation (orange), cumulative $N_2$ accumulation (black).

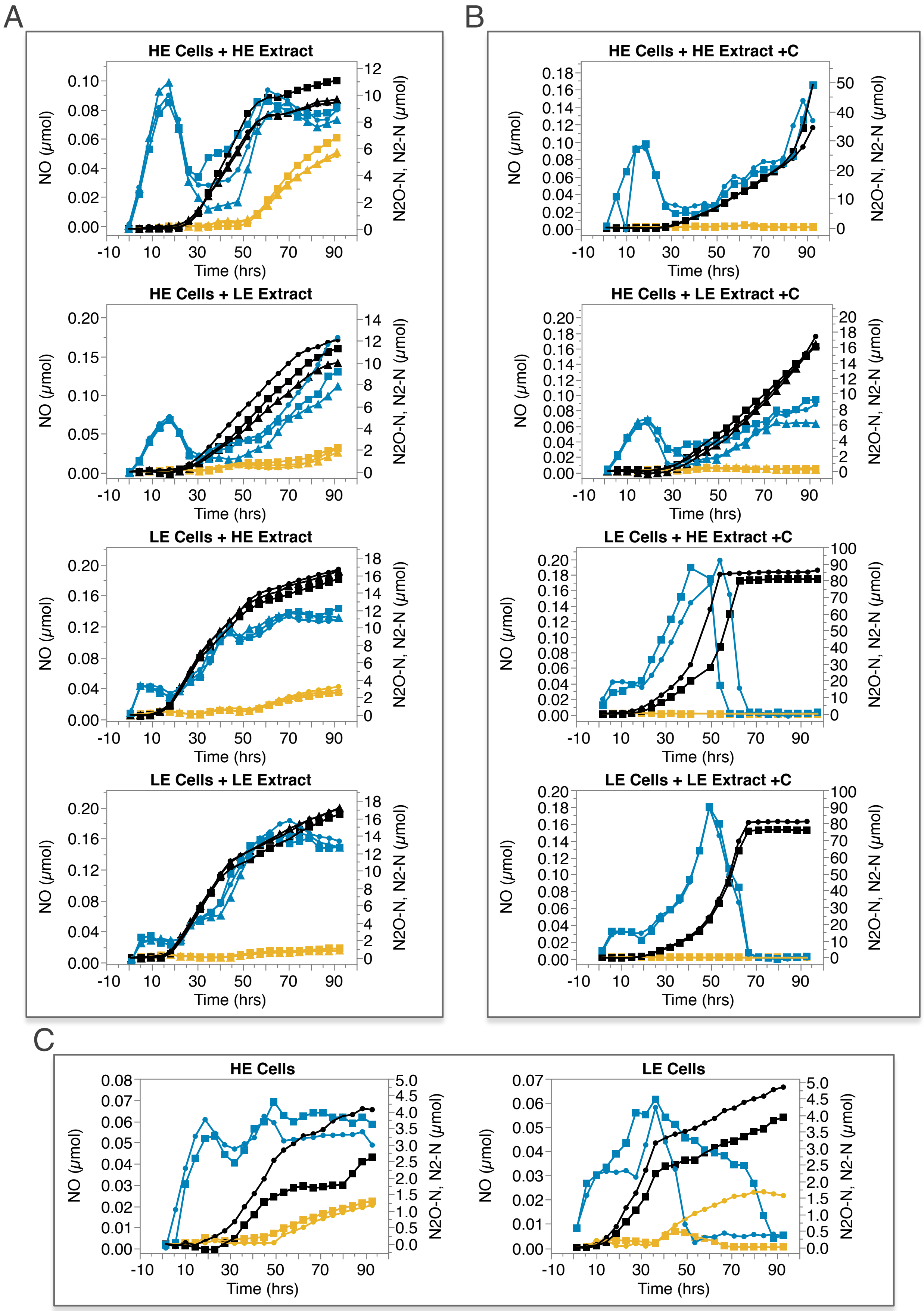
A
HE Cells + HE Extract
HE Cells + LE Extract
LE Cells + HE Extract
LE Cells + LE Extract
B
HE Cells + HE Extract +C
HE Cells + LE Extract +C
LE Cells + HE Extract +C
LE Cells + LE Extract +C
C
HE Cells
LE Cells
NO (µmol)
N2O-N, N2-N (µmol)
Time (hrs)

**Figure S3.** Inter-treatment variation in CBA-int denitrification gas emission kinetics and minimal $N_2O$ accumulation in glutamate amended treatments. Standard treatments (A), 3mM Glutamate amended controls (B), carbon negative controls (C). Headspace gases NO (blue), $N_2O$ (orange), $N_2$ (black) were quantified every 4hrs from triplicate (dots, squares, triangles) 3mM $NH_4NO_3$ amended anoxic extracted cell and chemistry based incubations. Control treatments (B, C) were carried out in minimum duplicate vials. Note separate scales between treatments to highlight relative gas accumulation.

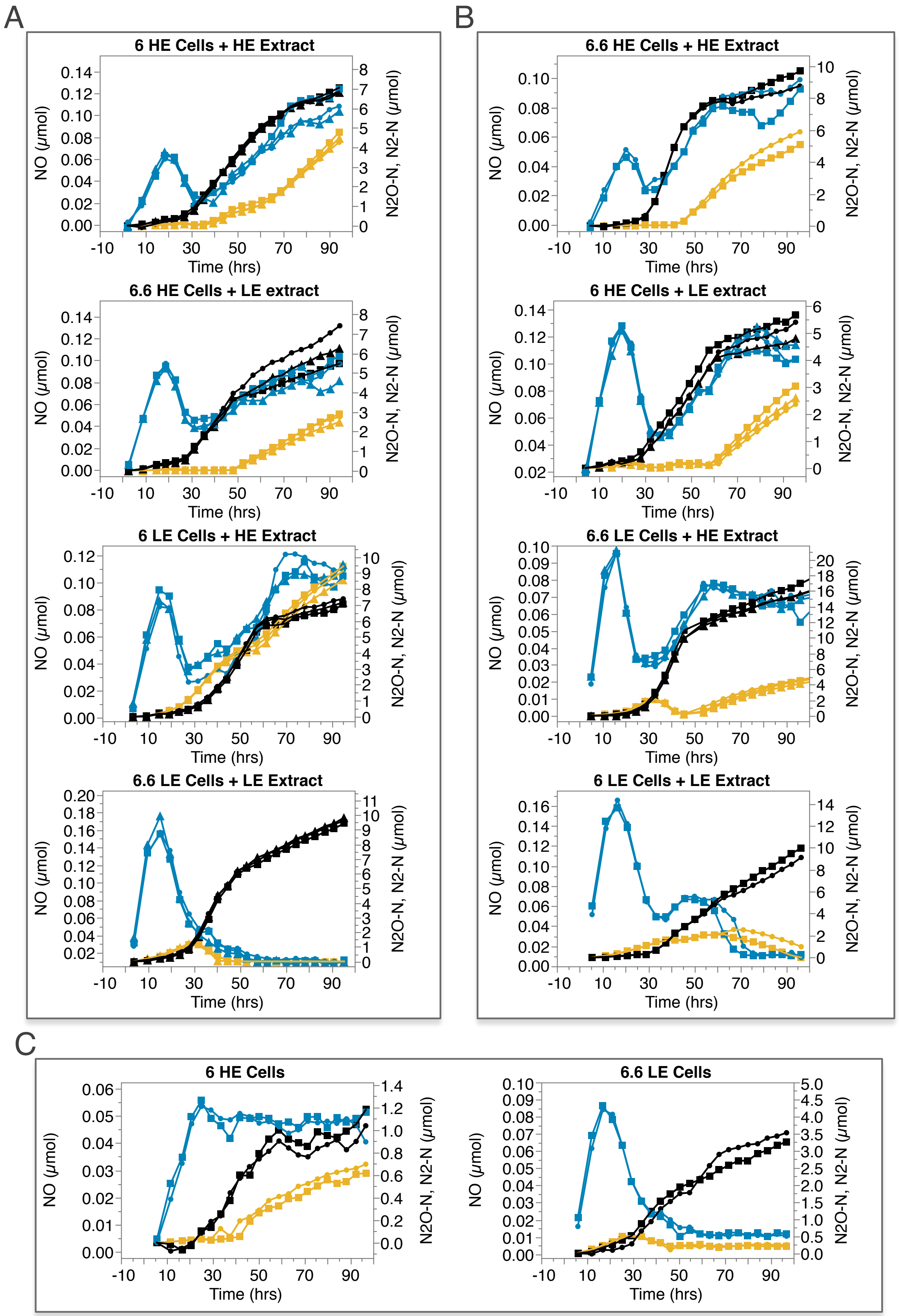
A
6 HE Cells + HE Extract
6.6 HE Cells + LE extract
6 LE Cells + HE Extract
6.6 LE Cells + LE Extract
B
6.6 HE Cells + HE Extract
6 HE Cells + LE extract
6.6 LE Cells + HE Extract
6 LE Cells + LE Extract
C
6 HE Cells
6.6 LE Cells
NO (μmol)
N2O-N, N2-N (μmol)
Time (hrs)

**Figure S4**. Contrasting denitrification gas emission kinetics between CBA-pH treatments derived from parent soils with contrasting pH and $N_2O$ emission potential. Standard treatments (A), alternative pH controls (B), carbon negative controls (C). Headspace gases NO (blue), $N_2O$ (orange), $N_2$ (black) were quantified every 4hrs from triplicate (dots, squares, triangles) 3mM $NH_4NO_3$ amended anoxic extracted cell and chemistry based incubations. Control treatments (B, C) were carried out in minimum duplicate vials. Chemical extract media pH reflected the contrasting pH of the parent soils (HEE: 6, LEE: 6.6) in standard treatments whereas pH and chemical extract were decoupled in alternative pH controls. Note separate scales between treatments to highlight relative gas accumulation.

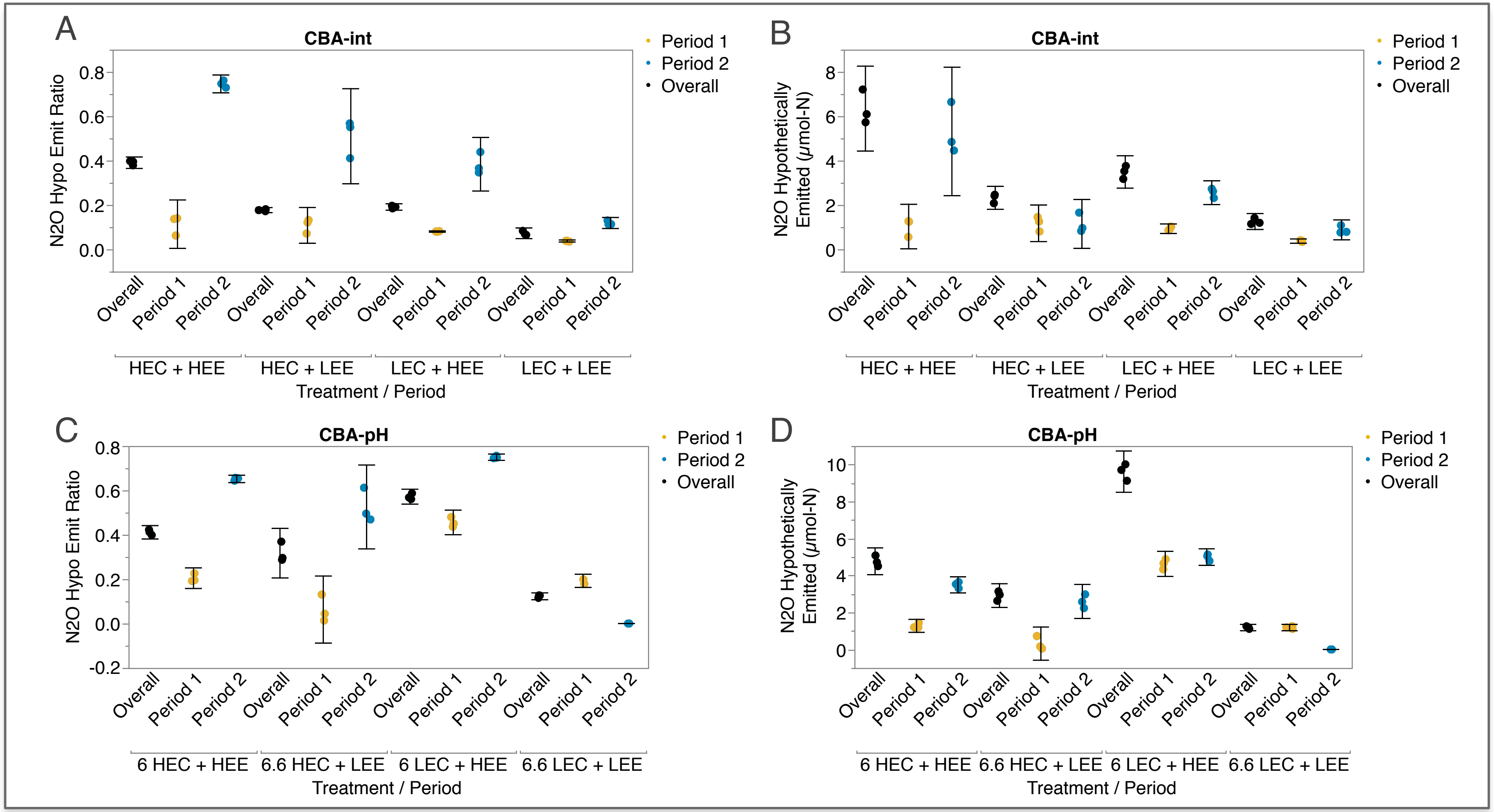


**Figure S5.** Comparison of cell based assay $N_2O$ ratios and $N_2O$ emitted (µmol-N per vial) across different assay periods. CBA-int (A,B) or CBA-pH (C,D). Cell based assay gas accumulation kinetics were split into an overall period (black), period 1 (orange) and period 2 (blue). The cutoff between period 1 and 2 was the point at which $N_2$ production rates decreased below, and consistently stayed below, 60% max $N_2$ accumulation rate for a given incubation. $N_2O$ ratios and $N_2O$ emitted summarise the $N_2O$ emission potential from CBA anoxic incubations amended with 3mM $NH_4NO_3$ and are calculated as $N_2O/N_2O+N_2$ and total $N_2O$ accumulated at the end of a defined incubation period, where phases of net negative $N_2O$ accumulation are ignored to account for multiple gas peaks. Triplicate vials were included per treatment. Results from triplicate vials per treatment are presented with 95% confidence intervals.

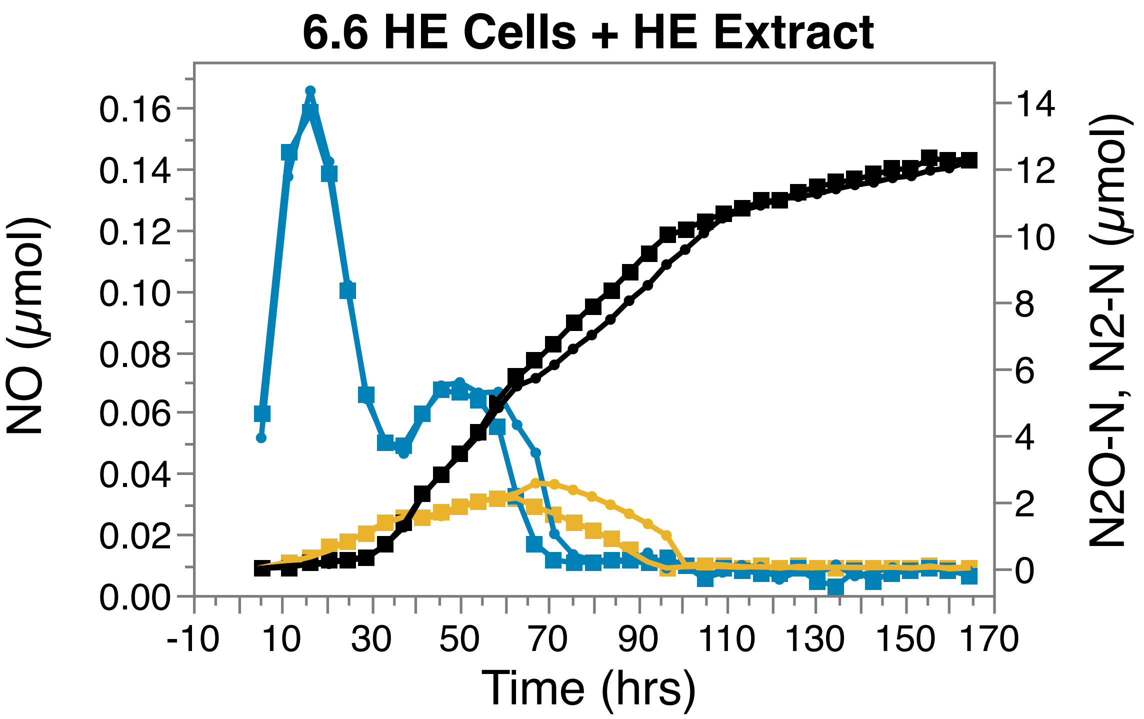


**Figure S6.** Extended 6 HEC + LEE CBA-pH incubation highlights additional $N_2$ linear rate switches. Headspace gases NO (blue), $N_2O$ (orange), $N_2$ (black) were quantified every 4hrs from duplicate (dots, squares, triangles) 3mM $NH_4NO_3$ amended anoxic extracted cell and chemistry based incubations.

**Supplementary Document S1**

### Partitioning of $N_2O$ emissions into two periods

Treatments varied greatly in their early vs. late $N_2O$ accumulation responses. To account for this we partitioned kinetics profiles into two distinct periods: period 1 prior to, and period 2 after, a sudden drop in $N_2$ accumulation rate (defined as a reduction in $N_2$ rate below 60% max rate for the same treatment).

Period 2 $N_2O$ ratio and emissions were typically larger than period 1 (Figure S5**Error! Reference source not found.**). Further, period 2 treatment rankings emulated the trends of the overall analyses across both assays, suggesting they determined the overall emission pattern. Though reduced confidence in period 2 treatments differences should be noted due to increasing occurrence of overlapping 95% confidence intervals. Period 1 $N_2O$ ratio and emissions did not emulate the overall trends due to similar $N_2O$ ratio and emissions for all treatments in the CBA-int (Figure S5A, B), and the alternative timing of $N_2O$ accumulation between treatments in CBA-pH (Figure S5C, D, FigureS4A). The relative impact of chemical extracts on $N_2O$ emission potential increased in period 2, becoming the most important determinant of $N_2O$ emissions in CBA-int and both $N_2O$ ratio and emissions in CBA-pH (Table S4, S5).
